# Distinct prefrontal projections conversely orchestrate social competition and hierarchy

**DOI:** 10.1101/2023.01.04.522808

**Authors:** Tae-Yong Choi, Hyoungseok Jeon, Sejin Jeong, Eum Ji Kim, Jeongseop Kim, Yun Ha Jeong, Byungsoo Kang, Murim Choi, Ja Wook Koo

## Abstract

Social animals compete for limited resources, resulting in a social hierarchy. Although different neuronal subpopulations in the medial prefrontal cortex (mPFC), a hub brain region of social hierarchy, encode distinct social competition behaviors, their identities and associated molecular underpinnings have not yet been identified. In this study, we found that mPFC neurons projecting to the nucleus accumbens (mPFC-NAc) encode social winning behavior, whereas mPFC neurons projecting to the ventral tegmental area (mPFC-VTA) encode social losing behavior by monitoring and manipulating neural circuit activity. High-throughput single-cell transcriptomic analysis and projection-specific genetic manipulation revealed that the expression level of POU domain, class 3, transcription factor 1 (*Pou3f1*) in mPFC-VTA neurons controls social hierarchy. Optogenetic activation of mPFC-VTA neurons increases *Pou3f1* expression and lowers social rank. Together, these data demonstrate that discrete activity and gene expression in separate mPFC projections oppositely orchestrate social competition and hierarchy.

## Introduction

Many social animals compete for limited resources such as food and territory, resulting in a social hierarchy within a group. An individual’s social rank determines the behaviors that are appropriate for that position^1,2,3,4,5,6,7,8^. For instance, social subordinates avoid competing with social dominants to prevent physical harm and energy loss, whereas social dominants engage in agonistic interactions to obtain easy access to food and sexual partners. Therefore, social status-specific behavioral adaptations are advantageous for individuals’ health and survival in an evolutionary context^9,10,11,12^.

The medial prefrontal cortex (mPFC) is an important hub region of social competition and hierarchy^13,14,15,16,17,18,19^. The mPFC neurons are activated by effortful behaviors intended for winning, such as push and retreat during the social dominance tube test^20^. Modifying the synaptic efficacy or neuronal activity in the mPFC regulates social dominance^20,21,22,23^. Recent studies have revealed that social losing (i.e., retreat) and winning behaviors during the tube test activate distinct mPFC neuronal subpopulations, suggesting that they separately encode opposite aspects of social competition and hierarchy^24,25^.

Although mPFC neurons send their axonal projections to various brain regions, it has been found that only hypothalamic neural pathways from the mPFC have been associated with social competition behaviors. Evidence shows that the mPFC neuronal subpopulations that give projections to different hypothalamic regions separately involve distinct aspects of agnostic behaviors^26^. These data support the idea that the mPFC controls specific behavioral features of social competition at the circuit level. Moreover, the mPFC neuronal subpopulations defined by connectivity have projection-specific molecular features^27,28,29,30^. Together, these results suggest that the mPFC may orchestrate social competition behaviors by manipulating gene expression and neuronal activity at the circuit level. Although molecular candidates in the mPFC that are involved in social dominance have been identified at the brain-region level^31,32^, the genes differentially expressed according to social status and their functions at the level of mPFC circuits remain unclear.

To address these questions, we identified differently activated mPFC projections in winner and loser mice using the tube test. We found that mPFC neurons projecting to the nucleus accumbens (mPFC-NAc) or ventral tegmental area (mPFC-VTA), which were previously reported to play an opposite role in social competition^33,34,35,36^, were activated in winners or losers, respectively. These observations were validated by measuring and manipulating the activities of these two mPFC projections during social competition. Single-cell RNA sequencing (scRNA-seq) of the mPFC from social dominants and subordinates was performed to investigate the projection-specific transcriptome of the mPFC-NAc or mPFC-VTA neurons. The subsequent projection-specific validation and genetic manipulation revealed that POU domain, class 3, transcription factor 1 (*Pou3f1*) in mPFC-VTA neurons was identified as a crucial factor in social hierarchy determination. Taken together, these results reveal that different neural circuit activities and gene expression in separate mPFC projections conversely control social competition and hierarchy.

## Results

### Social competition activates the mPFC-NAc or mPFC-VTA in the winner or loser, respectively

To investigate whether separate mPFC projections are differentially activated in social winners or losers in the tube test, we traced activated mPFC neurons and their projections using a Targeted Recombination in Active Population under Fos promoter (FosTRAP) strategy^37,38,39,40^. Two viral vectors were injected into the mPFC of *Fos-creER^T^*^2^ mice: AAV5-hSyn-mCherry to label all infected mPFC neurons and their projections and AAV5-EF1a-DIO-EYFP to label activated mPFC neurons and their projections (Figure 1A). A tube test was performed with several pairs of mice to determine the winner and loser 24 h after tamoxifen injection (Figure 1B). The resultant numbers of activated mPFC neurons in winners and losers were not different, but both were higher than those in the control group (Figure 1C, D). However, the NAc of the winner and the VTA of the loser received more activated mPFC projections (Figure 1C, E). Other brain areas, such as the lateral hypothalamus (LH), mediodorsal thalamus (MDT), basolateral amygdala (BLA), and periaqueductal gray (PAG), received similar amounts of activated mPFC projections from winners and losers (Figure S1A, B). To clarify whether mPFC-NAc or mPFC-VTA projection was selectively activated in winners or losers, we conducted Tracing Retrogradely the Activated Cell Ensemble (TRACE) experiments^41^. Two viral vectors that express Cre-inducible EYFP and mCherry retrogradely (rAAV2-retro (AAVrg)^42^) were injected into the NAc and VTA of *Fos-CreER^T^*^2^ mice, respectively. The animals were subjected to the tube test and subsequently administered 4-hydroxytamoxifen 2 h after the test (Figure 1F). Consequently, EYFP-expressing (i.e., NAc-projecting) and mCherry-expressing (i.e., VTA-projecting) mPFC neurons increased in winners and losers, respectively (Figure 1G, H). These results indicate that the mPFC-NAc projection is activated by social winning, whereas the mPFC-VTA projection is activated by social losing.

**Figure 1.**
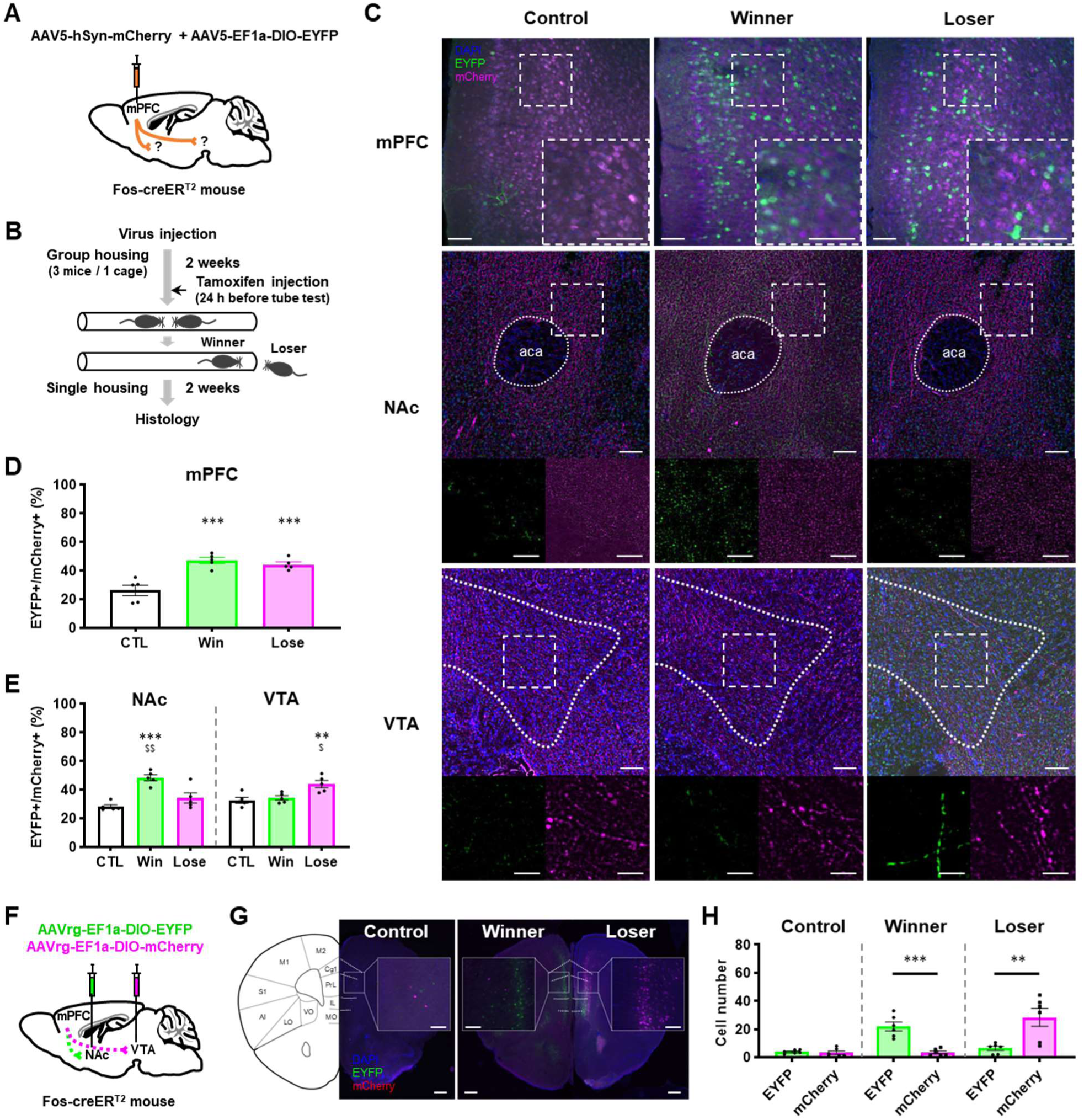
Social competition activates mPFC-NAc in winners and mPFC-VTA in losers. (A, B) Schematic illustration of viral injection (A) and behavioral test (B) for tracking activated mPFC neurons projecting to the NAc or VTA. (C) Representative images containing images of cell bodies in the mPFC (top) or images of projections in the NAc (middle) and VTA (bottom), and magnified images of the area indicated by a white dashed box. Scale bars, 0.1 mm (top); 0.01 mm (magnified images in middle and bottom). (D) Quantification of activated neurons by the ratio of EYFP^+^ (or FosTRAPed) cell numbers per mCherry^+^ (or infected) cell numbers in mPFC. n = 5 mice in each group; P < 0.001 by one-way ANOVA with Tukey’s test (CTL vs Win, P < 0.001***; CTL vs Lose, P < 0.001***; Win vs Lose, P = 0.6415). (E) Quantification of activated projections by the ratio of EYFP^+^ (or FosTRAPed) axon terminals per mCherry^+^ axon terminals (or total infected mPFC terminals) in ROIs. n = 5 mice in each group; P < 0.001 (NAc; CTL vs Win, P < 0.001***; CTL vs Lose, P = 0.2405; Win vs Lose, P = 0.0047^$$^), and P = 0.0044 (VTA; CTL vs Win, P = 0.8653; CTL vs Lose, P = 0.0059**; Win vs Lose, P = 0.0148^$^) by one-way ANOVA with Tukey’s test. (F) Schematic illustration of TRACE experiments. (G) Representative images containing the mPFC from control, winner, or loser mice, and magnified image of the area indicated by a white dashed box. Left illustration shows the coronal view of the mPFC. Scale bars, 0.5 mm; 0.1 mm (magnified images) (H) Quantification of EYFP^+^ (or mPFC-NAc) or mCherry^+^ (or mPFC-VTA) neurons in the mPFC from control, winner, or loser. n = 6 mice in each group; P < 0.001*** (Winner), P = 0.6596 (Control), and P = 0.0065** (Loser) by Student’s t-test (unpaired, two-tailed).

Next, we tested whether these two mPFC projections were activated in social dominants (rank 1, R1) or subordinates (rank 4, R4) after acquiring stable social ranks. Neuronal activity measured by c-fos expression (Figure S1C-E) or intrinsic excitability (Figure S2A, B, H, I) did not differ between R1 and R4 in either mPFC projection. c-Fos expression in the mPFC (Figure S1F) was similar between R1 and R4. Moreover, intrinsic electrophysiological properties such as rheobase current (Figure S2C, D, J, K), membrane capacitance (C_m_) (Figure S2E, L), resting input resistance (R_In_) (Figure S2F, M) and resting membrane potential (Figure S2G, N) did not differ between R1 and R4 in both mPFC projections. These results suggest that the activity of mPFC-NAc or mPFC-VTA circuitry is not involved in social dominance after social rank is formed. Furthermore, the structural connectivity of the mPFC-NAc or mPFC-VTA projection between social dominants and subordinates was not different (Figure S1G, H). It means that the differences in mPFC-NAc or mPFC-VTA circuit activity, but not anatomical changes, affect social competition.

### Manipulating the mPFC-NAc or mPFC-VTA activity conversely modulates social competition and hierarchy

Next, we investigated whether manipulating the activity of either the mPFC-NAc or mPFC-VTA projection would regulate social competition behaviors in an opposing manner. To examine the behavioral effects of mPFC-NAc or mPFC-VTA inhibition, a tube test was performed on multiple pairs of mice, each with a control and a cagemate mouse that had received viral vectors that were able to genetically ablate each projection-specific neuronal subpopulation: AAVrg-mCherry-IRES-Cre into the NAc or VTA to retrogradely express Cre recombinase and AAV2-Flex-taCasp3-TEVp into the mPFC to induce Cre-dependent apoptosis^43^ (Figure 2A, B, E, F). As a result, mice with ablating mPFC-NAc projection were significantly more prone to lose (Figure 2C) and exhibited fewer pushes and more retreats during the tube test (Figure 2D). Conversely, mice with inhibited mPFC-VTA projection displayed significantly more wins (Figure 2G), with a decrease in retreats and an increase in pushes (Figure 2H). We also performed a direct social interaction test and found no difference in the social interaction time between the control and circuit-inhibited mice (Figure S2O, P). These results suggest that the mPFC-NAc and mPFC-VTA circuitries conversely regulate social competition, which is not associated with sociability.

**Figure 2.**
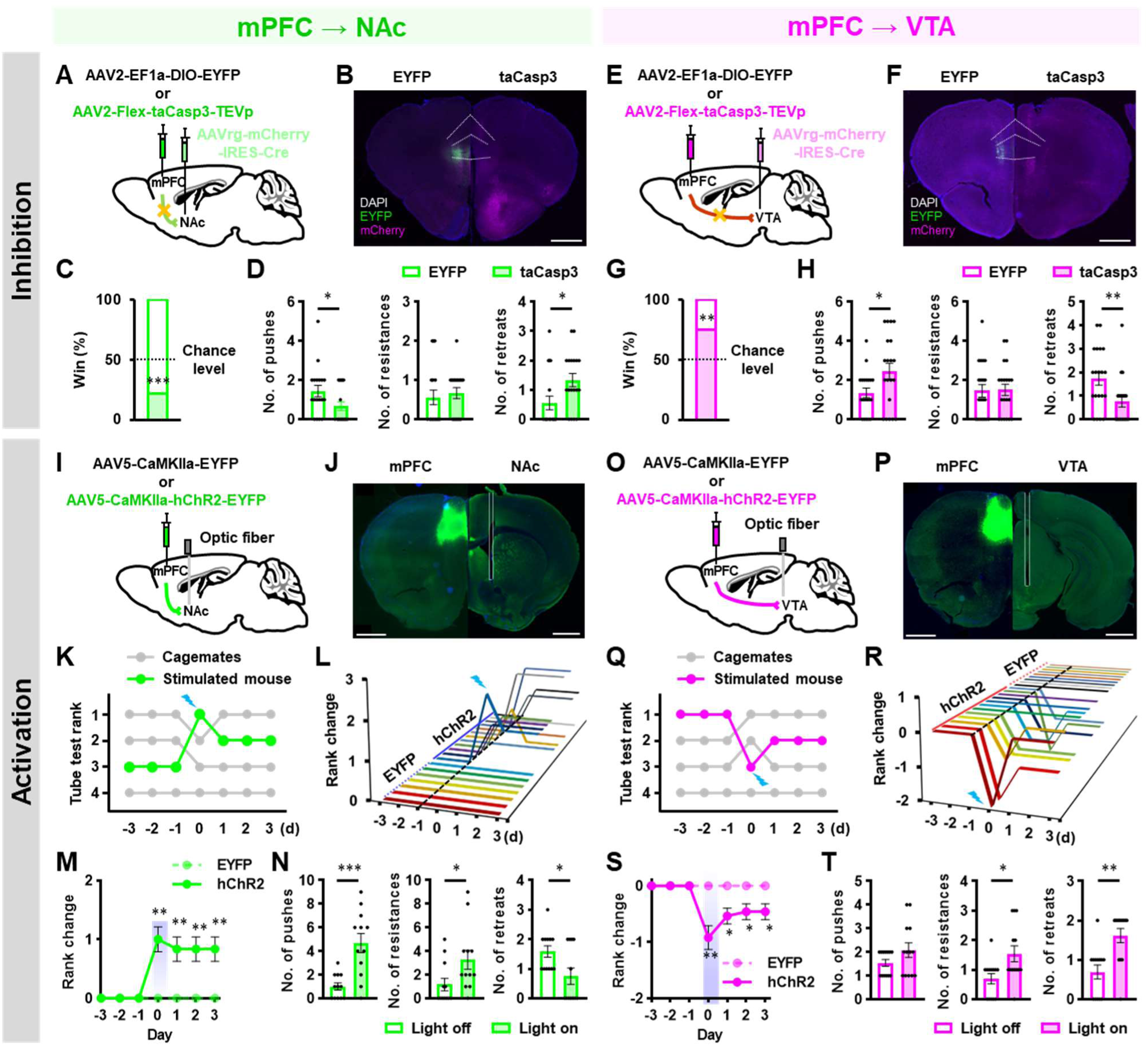
Manipulation of mPFC-NAc or mPFC-VTA activity oppositely modulates social competition and hierarchy. (A, E) Schematic illustration of genetic strategy to ablate mPFC-Nac (A) or mPFC-VTA (E) projection. (B, F) Representative images of control (left) or genetic ablation (right) in mPFE-NAc (B) or in mPFC-VTA (F) projection. Scale bar, 1 mm. (C, G) Genetic ablation of mPFC-NAc projection decreases (C), but mPFC-VTA projection increases (G) social dominance. n = 18 or 20 pairs for control and genetic ablation of mPFC-NAc (C) or mPFC-VTA (G), respectively; P <0.001*** (c, mPFC-NAc) and P = 0.0016** (G, mPFC-VTA) by Chi-square test (two-sided). (D, H) Behavioral performance of the control and mPFC-NAc-inhibited (D) or mPFC-VTA-inhibited (H) mice. n = 18 (D) or 20 (H) pairs of control and genetic ablation; P = 0.0403* (D, left, Push), P = 0.4763 (middle, Resistance), P = 0.0135* (right, Retreat), P = 0.0480* (H, left, Push), P = 0.8526 (middle, Resistance), and P = 0.0086** (right, Retreat) by Mann-Whitney U test (two-tailed). (I, O) Schematic illustration of optogenetic activation of mPFC-NAc (I) or mPFC-VTA (O) projection. (J, P) Representative images of the expression of hChR2 in mPFC (left) and optic fiber implantation in the NAc (J, right) or VTA (P, right) projections. Scale bar, 1 mm. (K, Q) Results of daily tube test conducted on a cage of four mice before and after acute photostimulation of mPFC-NAc projection of the rank 3 mouse (K) or mPFC-VTA projection of the rank 1 mouse (Q) on day 0. (L, R) Rank change following optogenetic stimulation of mPFC-NAc (L) or mPFC-VTA (R) projection in the tube test. One animal is shown on each line. On day 0, photostimulation was applied during the tube test. (M, S) Optogenetic activation of mPFC-NAc projection increases social ranks (M), but optogenetic activation of mPFC-VTA projection decreases social ranks (S). n = 7 cages for EYFP and n = 12 for hChR2 in mPFC-NAc (M), n = 8 for EYFP and n = 13 for hChR2 in mPFC-VTA (S); P = 0.0030** (M, 0 day), P = 0.0100** (M, 1-3 day), P = 0.0044** (S, 0 day), P = 0.0180* (S, 1 day), and P = 0.0456* (S, 2-3 day) by Mann-Whitney U test (two-tailed). (N, T) Behavioral performance of the same hChR2-expressing mice in mPFC-NAc (N) and mPFC-VTA (T) projections before and after photoactivation. Only mice showing rank changes are analyzed. n = 12 (N), n = 13 (T); P = 0.0010*** (N, left, Push), P = 0.0205* (middle, Resistance), P = 0.0415* (right, Retreat), P = 0.2828 (T, left, Push), P = 0.0206* (middle, Resistance), and P = 0.0019** (right, Retreat) by Mann-Whitney U test (two-tailed).

We then tested whether the activation of mPFC-NAc or mPFC-VTA projection changes social competition and hierarchy. A viral vector was administered to four cage mates to express channelrhodopsin proteins (hChR2) in the mPFC, and a fiber optic cannula was implanted into the NAc or VTA to deliver light stimulation (Figure 2I, J, O, P). After confirming stable social ranks, which were defined as the same ranking in the tube test for several consecutive days^20,23^, blue laser light was delivered to the mPFC-NAc or mPFC-VTA projection during the next tube test. The consequent photoactivation of mPFC-NAc projection in social subordinates increased their social rank (Figure 2K-M, S2Q, Video S1). The activation of mPFC-NAc projection specifically increased the number of pushes and resistances, while decreasing the number of retreats (Figure 2N). In contrast, the photoactivation of mPFC-VTA projection in social dominants decreased their social rank (Figure 2Q-S, S2R, Video S2). Furthermore, the activation of mPFC-VTA projection significantly increased the number of retreats (Figure 2T). Together, these results imply that the mPFC-NAc or mPFC-VTA projection conversely regulates social competition and hierarchy.

### Different social competition behaviors are separately encoded in the mPFC-NAc and mPFC-VTA

To elucidate when and how the mPFC-NAc or mPFC-VTA projection was activated during social competition, we measured the activity of these two mPFC projections during the tube test using fiber photometry (Figure 3A-C). AAVrg-CAG-Cre was injected into the NAc or VTA to retrogradely express Cre recombinase, and AAVdj-EF1a-DIO-GCaMP6s was injected into the mPFC. A fiber optic cannula was then implanted above the viral injection site of the mPFC to measure neuronal population activity. We found that the mPFC-NAc neurons were activated when a mouse pushed (Figure 3D, E, L) or approached opponents (Figure S3A, B, Q). The mPFC-NAc neurons were also activated when a mouse resisted opponent’s pushes (Figure S3C, D, R). However, the activity of mPFC-NAc neurons did not change when an animal retreated during the tube test (Figure 3F, G, M). These results suggest that the mPFC-NAc neurons encode active behaviors to win (i.e., push and approach) or to not lose (i.e., resistance).

**Figure 3.**
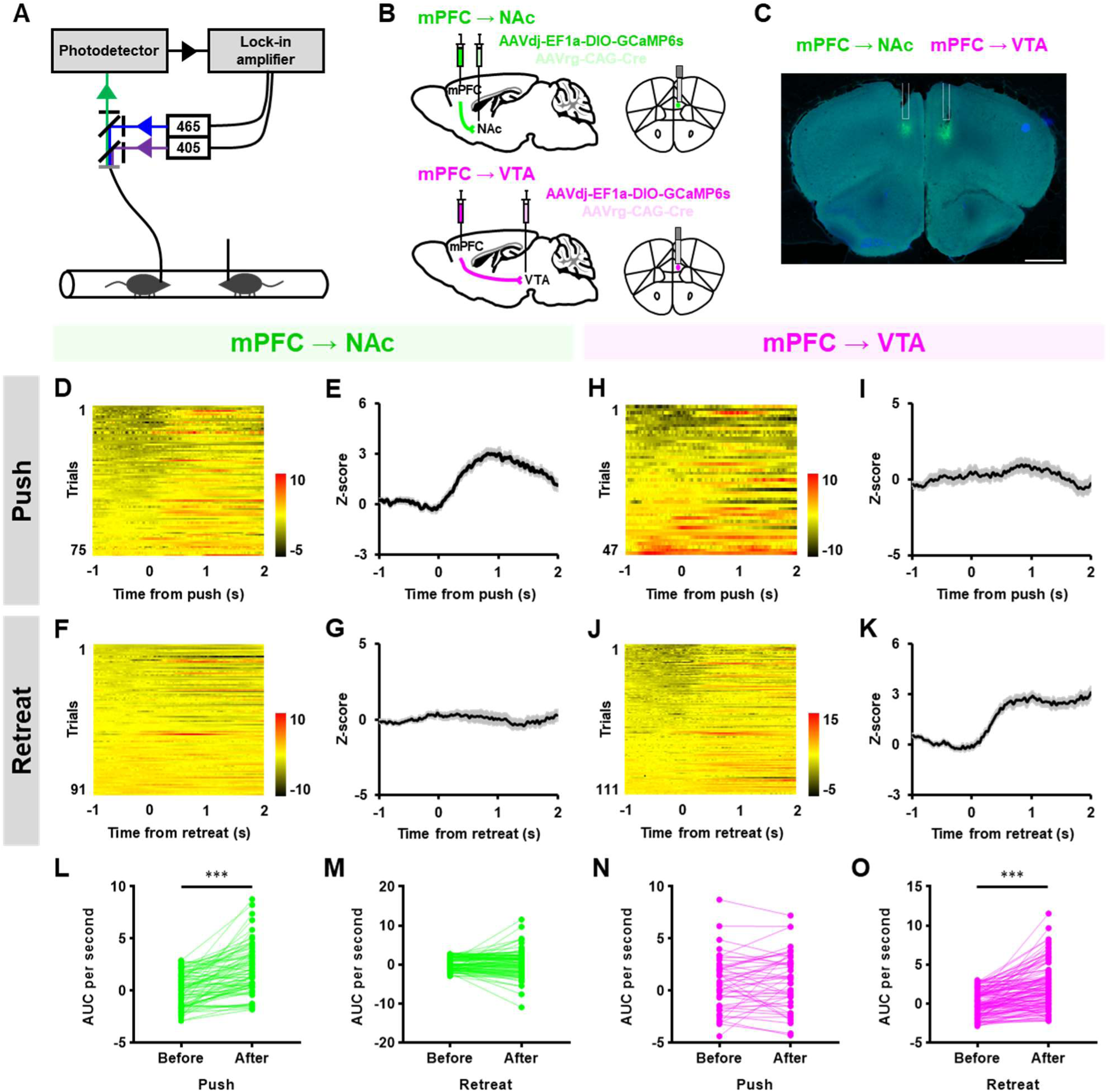
Different social competition behaviors are separately encoded in mPFC-NAc or mPFC-VTA neurons. (A-B) Schematic illustration of fiber photometry during the tube test (A) and virus injection and optic fiber implantation (B). (C) Representative images of a recording site and GCaMP6s expression of mPFC-NAc (left) or mPFC-VTA neurons (right). Scale bar, 1 mm. (D-G) Heatmap (D, F) and peri-event plot (E, G) of Z-scored mPFC-NAc activity aligned to the onset of pushes (D, E) or retreats (F, G). n = 75 push and n = 31 retreat trials from 10 mice. (H-K) Heatmap (H, J) and peri-event plot (I, K) of Z-scored mPFC-VTA activity aligned to the onset of pushes (H, I) or retreats (J, K). n = 47 push and n = 111 retreat trials from 10 mice. (L-O) Quantification of the area under curve (AUC) per second before and after pushes (L, N) or retreats (M, O) in mPFC-NAc (L, M) or mPFC-VTA (N, O) neurons. n = 75 trials (L), n = 91 (M), n = 47 (N), and n = 111 (O); P < 0.001*** (L, O) and P = 0.2074 (M) by Wilcoxon matched-pairs signed rank test (two-tailed), P = 0.3003 (N) by Student’s t-test (paired, two-tailed).

Next, we determined that the mPFC-VTA neurons were activated when a mouse retreated during the tube test (Figure 3J, K, O). However, other social competition behaviors, such as pushes (Figure 3H, I, N), approaches (Figure S3I, J, S), and resists (Figure S3K, L, T), did not change the activity of mPFC-VTA neurons. These results indicate that social losing behavior, such as retreat, is encoded in mPFC-VTA neurons.

Retreat behavior is classified into two types based on its spontaneity: active retreat, which is a retreat behavior without the opponent’s pushes or approaches, and passive retreat, which is a retreat behavior triggered by an opponent’s pushes. Similar to a previous study^44^, active retreats accounted for the majority of the overall retreats (180/205 trials) in our study (Figure S3E-H, M-P). We then investigated whether active or passive retreat had a distinct effect on the activity of these two mPFC projections. Neither active nor passive retreats altered the activity of mPFC-NAc neurons (Figure S3E-H, U-W). Although retreat behaviors activated mPFC-VTA neurons (Figure 3J, K, O), there was no difference between active and passive retreats (Figure S3M-P, X-Z). These findings demonstrate that all retreat behaviors, regardless of spontaneity, activate the mPFC-VTA neurons.

### Single-cell transcriptomic analysis reveals projection-specific molecular candidates for social hierarchy

To investigate the projection-specific molecular determinants in the mPFC-NAc or mPFC-VTA neurons participating in social hierarchy, we conducted single-cell RNA sequencing (scRNA-seq) of the mPFC of social dominants (R1) and subordinates (R4). The mPFC tissue punches were collected and dissected, and the resulting cells were subjected to a 10X Genomics scRNA-seq pipeline (Figure 4A). After excluding low-quality and doublet cells, resulting 41,409 cells (15,272 cells from R1 and 26,137 from R4) were clustered and analyzed (Figure S4A, Table S1-2). Of these, 14,883 cells (5,325 cells from R1 and 9,558 from R4) were classified as neurons (Figure 4B, Figure S4B) and were analyzed separately. Subsequently, we identified mPFC-NAc and mPFC-VTA neurons using previously reported projection-specific marker genes^28^. A principal component analysis plot showed a separate expression pattern of mPFC-NAc or mPFC-VTA projection-specific marker genes (Figure S4C-E). In the uniform manifold approximation and projection plot, mPFC-NAc projection marker genes were consistently enriched in cluster 1 (C1) and C8, and mPFC-VTA projection marker genes were higher in C0 and C3 (Figure 4C). Thus, we assumed C1 and C8 to be mPFC-NAc neurons and C0 and C3 to be mPFC-VTA neurons.

**Figure 4.**
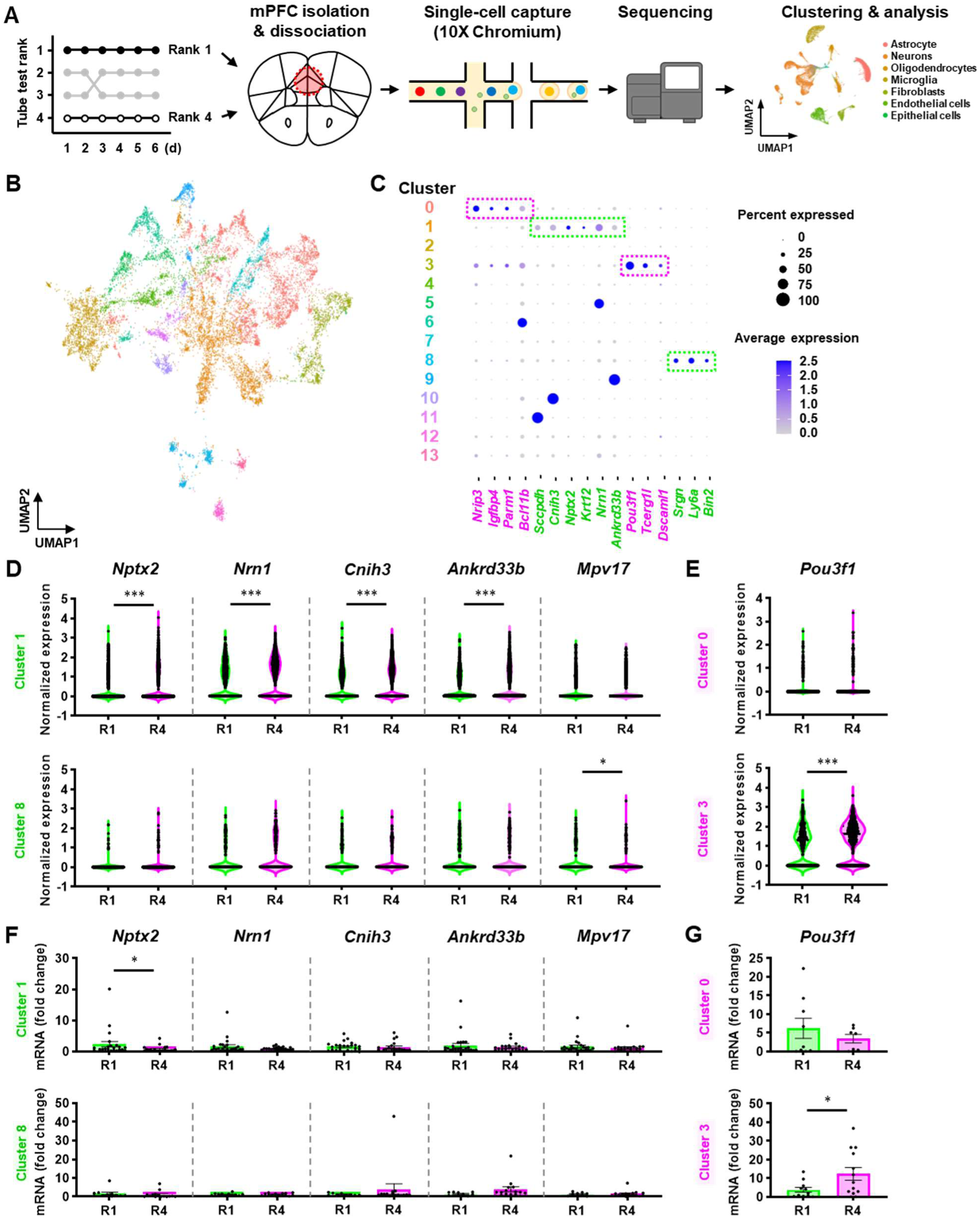
Single cell transcriptomic analysis reveals projection-specific candidate genes that regulate social hierarchy. (A) Schematic illustration of scRNA-seq of the mPFC from R1 and R4 mice. (B) A uniform manifold approximation and projection (UMAP) plot of 5,325 and 9,558 neurons from the mPFC of R1 (n =11) and R4 mice (n =11), respectively, using mPFC marker genes as defined by Kim et al^28^. (C) Expression of the projection-specific marker genes across identified clusters, with the colors of the clusters the same as in the UMAP in B. mPFC-NAc neurons, C1 and C8; mPFC-VTA neurons, C0 and C3. (D, E) DEGs between R1 and R4 in mPFC-NAc (D) or mPFC-VTA (E) neurons by scRNA-seq analysis. n = 1,331 neurons (D, C1, R1), n = 2,100 (D, C1, R4), n = 219 (D, C8, R1), n = 308 (D, C8, R4), n = 1,346 (E, C0, R1), n = 2,361 (E, C0, R4), n = 449 (E, C3, R1), and n = 675 (E, C3, R4); P < 0.001*** (*Nptx2, Nrn1, Cnih3, Ankrd33b* in D, C1; *Pou3f1* in E, C3), P = 1.0000 (*Mpv17* inD,C1), P = 1.0000 (*Nptx2*, *Cnih3, Ankrd33b* in D, C8; *Pou3f1* in E, C0), P = 0.3348 (*Nrn1* in D, C8), and P= 0.0451* (*Mpv17* in D, C8) by Wilcoxon Rank-Sum test. (F, G) scRT-qPCR validation of DEGs between R1 and R4 in mPFC-NAc (F) or mPFC-VTA (G) neurons. n = 24-25 neurons from 4 mice (F, C1, R1), n = 21, 4 (F, C1, R4), n = 11, 4 (F, C8, R1), n = 15, 4 (F, C8, R4), n = 9, 3 (G, C0, R1), n = 7, 3 (G,C0, R4), n =11, 3 (G, C3, R1), and n =13, 3 (G, C3, R4). P = 0.0177* (*Nptx2* in F, C1), P = 0.6581 (*Nrn1* in F, C1), P = 0.6661 (*Cnih3* in F, C1), P= 0.7890 (*Ankrd33b* in F, C1), P = 0.4806 (*Mpv17* in F, C1), P = 0.0751 (*Nptx2* in F, C8), P = 0.8722 (*Cnih3* in F, C8), P = 0.1085 (*Ankrd33b* in F, C8), P = 0.9884 (*Mpv17* in F, C8), and P = 0.9182 (*Pou3f1* in G, C0) by Mann-Whitney U test (two-tailed), P = 0.8371 (*Nrn1* in F, C8) and P = 0.0392* (*Pou3f1* in G, C3) by Student’s t-test (unpaired, two-tailed).

Next, we analyzed differentially expressed genes (DEGs) between R1 and R4 in each mPFC projection and found that the expression of *Pou3f1* was higher in R4 than in R1 in C3, but not in C0, of mPFC-VTA neurons (Figure 4E). To verify whether this gene is highly expressed in C3 of mPFC-VTA neurons in social subordinates compared to its expression in dominants, we measured the mRNA levels of *Pou3f1* and cluster-specific marker genes, such as *Nrip3* for C0, by single-cell reverse transcriptase quantitative polymerase chain reaction (scRT-qPCR) with aspirated intracellular contents of retrogradely labeled mPFC-VTA neurons from R1 or R4 mice (Figure S4F-H). The expression of *Pou3f1* mRNA in C3, which is determined by the significantly lower expression of *Nrip3,* but not that in C0, was higher in R4 than in R1 mice (Figure 4G). However, the expression level of *Pou3f1* in mPFC-VTA neurons (C0 + C3) was not different between R1 and R4 mice (Figure S4J). Moreover, *Nptx2*, *Nrn1*, *Cnih3*, and *Ankrd33b* in C1 and *Mpv* in C8 of the mPFC-NAc neurons were more highly expressed in social subordinates than they were in dominants (Figure 4D). However, the statistical differences calculated from scRNA-seq analysis were not validated by the scRT-qPCR results (Figure 4F). The expression levels of these genes in mPFC-NAc neurons (C1 + C8) were also not significantly different between R1 and R4 (Figure S4I). Together, these results suggest that the expression level of *Pou3f1* in mPFC-VTA neurons, especially in C3, may affect social hierarchy.

Although *Pou3f1* controls neurodevelopment and myelination^45^, it remains unclear how this gene in the mPFC-VTA neurons modulates the social hierarchy. Identifying the downstream genes regulated by the Pou3f1 transcription factor that binds to the octamer motif (5’-ATTTGCAT-3’) would thus be helpful^46,47,48,49^. Therefore, we re-analyzed previous chromatin immunoprecipitation sequencing (ChIP-seq) studies using Pou3f1^49,50,51^. Gene ontology (GO) enrichment analysis of genes near the Pou3f1 binding motifs revealed several notable terms on synaptic function (i.e., glutamatergic synapse, synaptic membrane, presynapse, etc.) as well as terms on neurodevelopment (i.e., forebrain development and positive regulation of nervous system development) (Figure S4K-M). This suggests that *Pou3f1* expression in mPFC-VTA neurons may control social competition and hierarchy by regulating synaptic functions in the neural circuit.

### Manipulating Pou3f1 expression in the mPFC-VTA regulates social hierarchy

To examine whether the expression level of *Pou3f1* in mPFC-VTA neurons regulates individuals’ social status, we first suppressed the expression of *Pou3f1* in mPFC-VTA neurons in social subordinates (i.e., R3 or R4) by expressing shRNA against *Pou3f1* (Figure 5A, B, S5A, B). Several weeks after the viral injection, the social rank of the social subordinates that received *Pou3f1* shRNA in mPFC-VTA neurons increased (Figure 5C-E, S5I). Similarly, the knockdown of *Pou3f1* in mPFC-VTA neurons of social subordinates reduced retreats and increased pushes (Figure 5F). To test whether the winner effect of the *Pou3f1* knockdown in mPFC-VTA neurons observed in the tube test can be transferred to another social dominance behavior, we developed a behavioral test named the wet-bedding avoidance (WBA) test, which permits the investigation of individuals’ territorial instincts to occupy the platform for avoiding wet bedding in a cage (Figure S5E, Video S3). We found that social ranks in the WBA test, which were determined by how long each mouse occupied the platform, were well correlated with the tube test ranks (Figure S5F-H). The total time on the platform and social ranks of the WBA test also increased after the knockdown of *Pou3f1* in mPFC-VTA neurons, similar to the tube test (Figure 5G).

**Figure 5.**
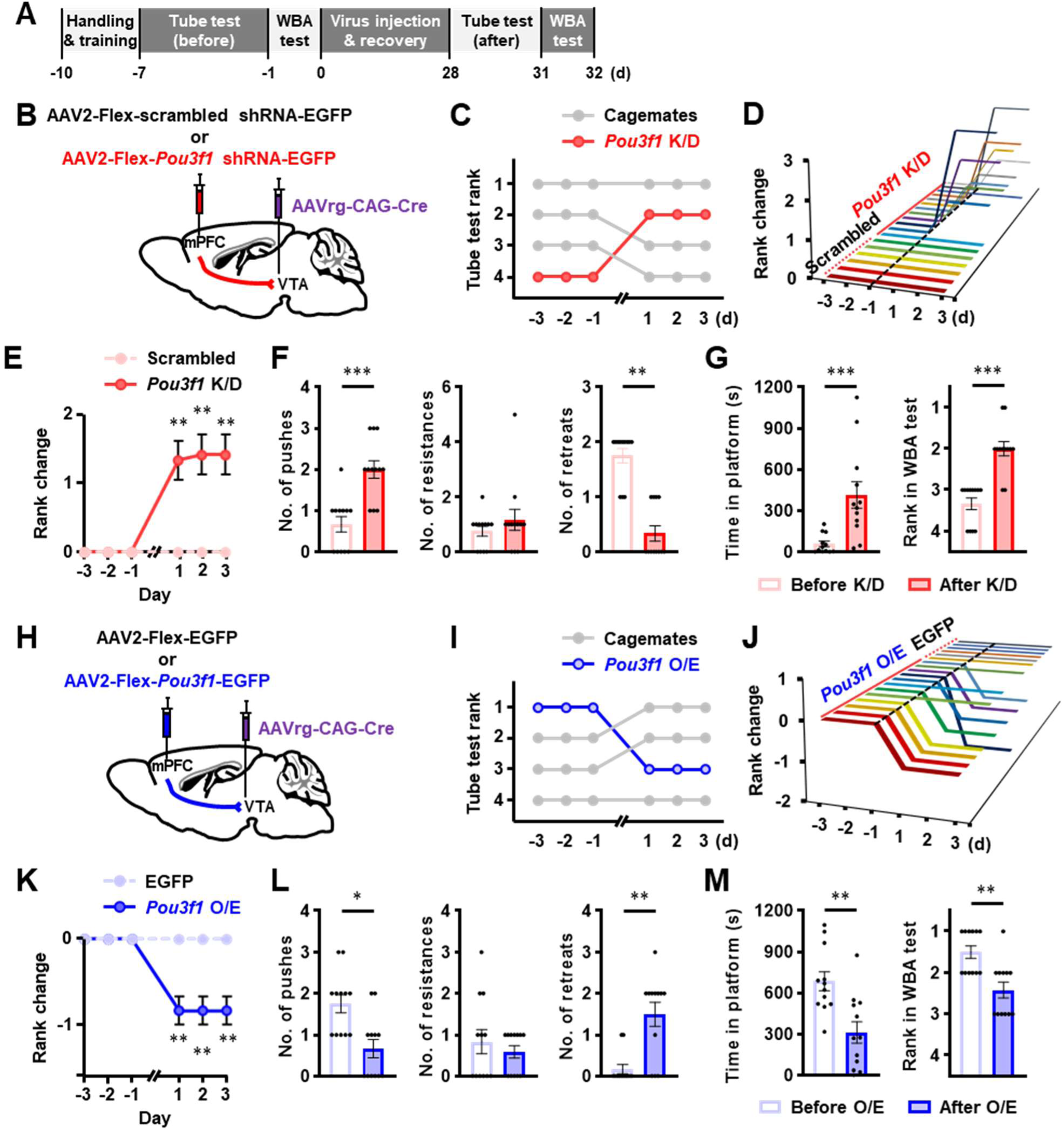
Manipulation of *Pou3f1* expression in mPFC-VTA neurons bidirectionally regulates social hierarchy. (A) Experimental schedule of behavioral tests for social competition and hierarchy before and after manipulating the expression of projection-specific DEGs. (B, H) Schematic illustration of the knock-down (B, *Pou3f1* K/D) or the over-expression (H, *Pou3f1* O/E) of *Pou3f1* in mPFC-VTA neurons. (C, I) Results of a daily tube test performed on a group of 4 mice before and after *Pou3f1* K/D in mPFC-VTA neurons of the R4 mouse (C) or *Pou3f1* O/E in mPFC-VTA neurons of the R1 mouse (I). (D, J) Rank change following *Pou3f1* K/D in mPFC-VTA neurons of social subordinates (D) or *Pou3f1* O/E in mPFC-VTA of social dominants (J) in the tube test. One animal is shown on each line. Viral vectors were injected on day 0. (E, K) *Pou3f1* K/D in mPFC-VTA neurons of social subordinates increases (E), but *Pou3f1* O/E in mPFC-VTA neurons of social dominants decreases their social ranks (K). n = 6 cages for scrambled shRNA and n =12 for *Pou3f1* K/D(E),n=6for EGFPandn=12for *Pou3f1* O/E in mPFC-VTA neurons (K); P = 0.0023** (E, 1 day), P = 0.0034** (E, 2-3 day), and P = 0.0075** (K, 1-3 day) by Mann-Whitney U test (two-tailed). (F, L) Comparison of behavioral performance of same mice before and after *Pou3f1* K/D (F) or O/E (L) in mPFC-VTA neurons. Only mice showing rank changes are analyzed. n = 12 cages (F, L); P = 0.0010*** (F, left, Push), P = 0.8125 (F, middle, Resistance), P = 0.0039** (F, right, Retreat), P = 0.0137* (L, left, Push), P = 0.5625 (middle, Resistance), and P = 0.0078** (right, Retreat). (G, M) Changes in time on the platform (left) and rank (right) in wet bedding avoidance (WBA) test by *Pou3f1* K/D (G) or O/E (M) in mPFC-VTA neurons. n = 12 cages (G, M); P < 0.001*** (G, Time in platform or Rank in WBA test), P = 0.0020** (M, Rank in WBA test) by Wilcoxon matched-pairs signed rank test (two-tailed), and P = 0.0023 (M, Time in platform) by Student’s t-test (paired, two-tailed).

Next, we next tested whether the overexpression of *Pou3f1* in mPFC-VTA neurons of social dominants (R1 or R2) decreased their social ranks (Figure 5A, H, S5C, D). The overexpression of *Pou3f1* in mPFC-VTA neurons decreased the tube test ranks (Figure 5I-K, Figure S5J) and the number of pushes, but increased the number of retreats during the tube test (Figure 5L). Additionally, the total amount of time occupying the platform and social ranks in the WBA test were reduced by the overexpression of *Pou3f1* in mPFC-VTA neurons (Figure 5M). These results indicate that the expression level of *Pou3f1* in mPFC-VTA neurons can modulate social competition and hierarchy.

### Activation of the mPFC-VTA increases Pou3f1 expression and reduces social dominance

We discovered that the optogenetic activation of mPFC-VTA projection or the overexpression of *Pou3f1* in mPFC-VTA neurons reduced social rank (Figure 2O-T, 5H-M). These findings raise the question of whether mPFC-VTA activity and *Pou3f1* expression are correlated with the regulation of the social hierarchy. To address this question, we directly stimulated mPFC-VTA neurons by injecting a retrograde viral vector to express Cre recombinase (AAVrg-CAG-Cre) into the VTA, as well as Creinducible control or hChR2-expressing viral vectors (AAV5-EF1a-DIO-EYFP or AAV5-EF1a-DIO-hChR2(H134R)-EYFP) into the mPFC to determine whether *Pou3f1* expression in these neurons was altered (Figure 6A). After confirming stable tube test ranks, optogenetic stimulation was applied to the social dominants (R1 or R2) during the tube test. As shown in Figure 2O-T, the photoactivation of mPFC-VTA neurons reduced the social rank (Figure 6B-D, Figure S6A). Three days after optogenetic stimulation, hChR2-expressing mPFC-VTA neurons were collected, and scRT-qPCR was performed to determine the expression level of *Pou3f1*. As shown in Figure 4G, *Pou3f1* expression in C3 was higher in social subordinates without optogenetic activation (R3/R4 + EYFP) than in social dominants without optogenetic activation (R1/R2 + EYFP) (Figure 6E). The photoactivation of mPFC-VTA neurons not only reduced the social rank but also increased the expression level of *Pou3f1* in C3 (Figure 6E). However, *Pou3f1* expression among the three groups was not different in C0 (Figure 6E) or whole mPFC-VTA neurons (C0 + C3) (Figure S6C). These results suggest that the activation of mPFC-VTA neurons increases the expression of *Pou3f1* in the C3 projecting subpopulation, thereby reducing the social hierarchy.

**Figure 6.**
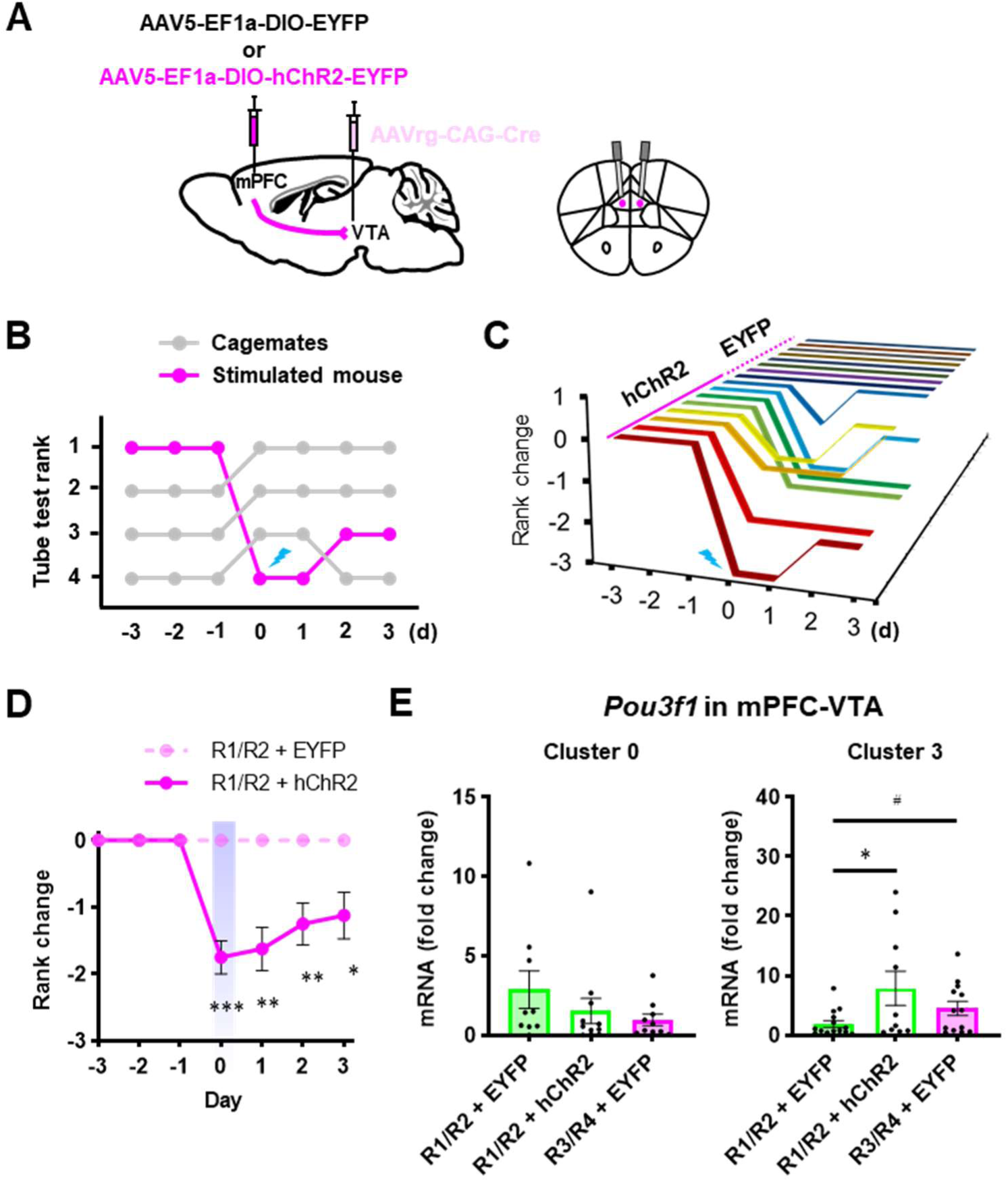
Optogenetic activation of mPFC-VTA neurons increases *Pou3f1* expression and decreases social ranks. (A) Schematic illustration of virus injection (left) and fiber optic implantation (right). (B) Results of a daily tube test conducted on a cage of 4 mice before and after soma-targeting optogenetic activation of mPFC-VTA neurons of the R1 mouse on day 0. (C) Rank change following optogenetic activation of mPFC-VTA neurons in the tube test. One animal is shown on each line. Photostimulation was applied during the tube test on day 0. (D) Optogenetic activation of mPFC-VTA neurons of social dominants decreases their social ranks. n = 8 cages for EYFP, and n = 8 for hChR2; P < 0.001*** (0 day), P = 0.0014** (1 day), P = 0.0070** (2 day), and P = 0.0256* (3 day) by Mann-Whitney U test (two-tailed). (E) *Pou3f1* expression level in mPFC-VTA neurons from social dominants (rank 1 or 2) without or with optogenetic activation of mPFC-VTA neurons or from social subordinates (rank 3 or 4). n = 9 cells from 4 mice (C0, R1/R2 + EYFP), n = 11, 4 (C0, R1/R2 +hChR2), n = 10, 4 (C0, R3/R4 + EYFP), n = 15, 4 (C3, R1/R2 + EYFP), n = 10, 4 (C3, R1/R2 +hChR2), n = 13, 4 (C3, R3/R4 + EYFP); P = 0.0345 by one-way ANOVA with Tukey’s test (C3, R1/R2 + EYFP vs R1/R2 + hChR2, P = 0.0265*; R1/R2 + EYFP vs R3/R4 + EYFP, P = 0.4150), and P = 0.0436^#^ (R1/R2 + EYFP vs R3/R4 + EYFP, Student’s t-test, unpaired, two-tailed).

## Discussion

In this study, we discovered that the mPFC-NAc and mPFC-VTA projections play opposing roles in social competition and hierarchy. This provides compelling evidence that distinct neuronal projections in the mPFC encode different social competition behaviors^20,21,24,25,26^. Furthermore, our projection-specific transcriptomic analysis with scRNA-seq data using mPFC projection-specific marker genes^28^ and subsequent genetic manipulations uncovered projection-specific molecular determinants for social dominance. Overall, these findings suggest that prefrontal functions in social dominance behaviors are distributed among highly specialized mPFC projections (e.g., mPFC-NAc vs. mPFC-VTA) that contain distinct molecular features or mechanisms for social competition and hierarchy (Figure S6D).

### Converse roles of mPFC-NAc and mPFC-VTA in social competition and hierarchy

The NAc and VTA are key components of the mesolimbic system, which controls reward and motivation by regulating dopaminergic neurotransmission from the VTA to NAc and other brain areas^52,53^. These two brain areas conversely control social hierarchy. Pharmacological lesions or inhibition of the NAc reduce the social dominance^34,35^, but pharmacological inhibition of the VTA increase the social dominance^33,36^. These results are consistent with the behavioral alterations that occur when the mPFC projections to these two brain areas are repressed (Figure 2A-H).

In the present study, one of our main findings is that the mPFC-NAc and mPFC-VTA projections conversely control social competition and hierarchy. This raises a question regarding whether these two mPFC projections operate independently or if one mPFC projection inhibits the other. We can rule out the possibility that the same mPFC neurons differentially control the NAc and VTA, because the mPFC-NAc and mPFC-VTA neurons are located in different layers (i.e., mPFC-NAc neurons in layer 2/3-5a and mPFC-VTA neurons in layer 5b-6) and few mPFC neurons project to both the NAc and VTA^27,28,30^. We found that pushing or retreating enhanced the activity of mPFC-NAc or mPFC-VTA neurons, respectively, but had no effect on the activity of another projection (Figure 3D-O). This implies that these two projections independently participate in social competition behaviors.

Additionally, we discovered that the mPFC-NAc neurons encode social winning behaviors such as push and resistance, while the mPFC-VTA neurons encode social losing behaviors such as retreat (Figure 3). These results strongly support previous findings that one-third of push-encoding mPFC neurons are also activated when a mouse resists an opponent’s pushes^20^ and retreat behavior is encoded in distinct neuronal subpopulations in the mPFC^24,25^. However, a discrepancy exists between our findings and those of previous study. Although other researchers have reported that distinct mPFC neurons encode push and approach^24,25^, our findings show that these two behaviors enhance the activity of mPFC-NAc neurons (Figure 3D, E, L, S3A, B, Q). This may be due to the restriction of fiber photometry, which measures the activity of neuronal populations without considering cellular heterogeneity. Different mPFC-NAc neuronal subpopulations may encode a push or an approach separately.

The mPFC neurons are connected with many other brain areas and can modulate social dominance through top-down circuitries by monitoring ongoing conflicts, evaluating different behavioral options, and contributing to behavioral responses^54,55,56^. In addition to the mPFC-NAc and mPFC-VTA projections in our study, the lateral hypothalamus-projecting mPFC neurons (mPFC-LH) also encodes social competition behaviors^21,26^. However, mPFC-LH projection was excluded in our study because it was not activated in either winners or losers in the tube test (Figure S1A, B). It suggests that different mPFC projections are involved depending on what animals compete for: mPFC-LH neurons encode social competition for food, whereas mPFC-NAc or mPFC-VTA neurons encode social competition for territory.

### Role of Pou3f1 in mPFC-VTA in social competition and hierarchy

*Pou3f1,* also known as *Oct6* (octamer-binding transcription factor 6), *Tst-1* (Testes-1), and *SCIP* (suppressed cyclic AMP (cAMP) inducible POU), was first identified in the embryonic neural tube and the adult brain^57^, as well as in myelin-forming glia^47^. Pou3f1 acts as a transcription factor that regulates the development of the nervous system in embryonic stem cells and the myelination of Schwann cells and oligodendrocytes^43,46,48,49,58^. *Pou3f1* is also an mPFC-VTA projection marker gene that is mostly found in cortical layer 5^28,30,59^. In our study, *Pou3f1* was identified as an mPFC-VTA projection-specific DEG between social dominants and subordinates. Social subordinates had greater expression level of *Pou3f1* than those of social dominants in the specific subpopulation of mPFC-VTA neurons, C3 (Figure 4E, G). Although it is technically difficult to manipulate *Pou3f1* expression in C3, we showed that manipulating the expression level of this gene in whole mPFC-VTA neurons regulates social competition and hierarchy (Figure 5). Moreover, the expression level of *Pou3f1* in C3 was decreased by the optogenetic activation of mPFC-VTA neurons (Figure 6). These findings provide compelling evidence that the activity of mPFC-VTA neurons alters *Pou3f1* expression, which, in turn, controls social dominance.

However, how the expression level of *Pou3f1* in mPFC-VTA neurons regulates social dominance behavior remains unclear. According to recent scRNA-seq studies using mPFC tissues, *Pou3f1* is a unique marker gene for a subtype of excitatory neurons in layer 5 of the mPFC^60,61,62,63^. Our GO analysis with Pou3f1 ChIP-seq data from previous literature revealed several notable terms, such as glutamatergic synapse, synaptic membrane and presynapse (Figure S4K-M). These suggest that *Pou3f1* expression in mPFC-VTA neurons controls social competition and hierarchy by regulating the excitatory synaptic function of the mPFC-VTA circuitry. Furthermore, *Pou3f1* expression can be increased by intracellular cAMP, a second messenger that is enhanced by the activation of G-protein coupled receptors coupled to Gs protein (Gs-GPCRs) and adenylyl cyclase^47^, or estrogen^65^. The activation of Gs-GPCRs raises intracellular cAMP levels, which in turn increases neuronal activity, and the deletion of a gene encoding the estrogen receptor β reduces aggressive behavior^65^. It suggests that Gs-GPCR-activating neuromodulators or estrogen can regulate social hierarchy by manipulating mPFC-VTA circuit activity and *Pou3f1* expression.

### Limitations of the study

Our study demonstrated that the mPFC orchestrates social competition behaviors by governing two discrete projections with opposite functions (i.e., mPFC-NAc vs. mPFC-VTA) as working units that perform specific behavioral features of social competition at the circuit level. As mentioned above, other mPFC projections may be involved in the regulation of social competition behaviors according to spontaneity (e.g., active vs. passive retreats), or reward attributes (e.g., food, territory, or mating). Additionally, the social competition can be controlled by certain neuronal ensembles in the mPFC at the cell type^23^ or circuit level^21^, which requires further study.

The projection-specific molecular/physiological underpinning of social competition behaviors in the mPFC also requires further investigated. The mechanism by which the neural activity of mPFC-VTA projections modulates *Pou3f1* expression and its related molecular/physiological mechanisms in mPFC-VTA neurons and *vice versa* has not been studied. Regarding the mPFC-NAc projection, the expression levels of *Nptx2*, *Nrn1*, *Cnih3*, and *Ankrd33b* in the C1, which were identified as DEGs in our scRNA-seq analysis but not corroborated by scRT-qPCR (Figure 4D, F), were highly associated with social ranks. In particular, these DEGs are projection marker genes, indicating that their molecular features are closely associated with circuit functions. A pivotal step toward a comprehensive understanding of the prefrontal functions underlying social competition behaviors would be to elucidate the behavioral/physiological implications of these projection-specific DEGs.

## Acknowledgements

We thank all members of behavioral neuroepigenetics laboratory in Korea Brain Research Institute (KBRI) for helpful discussion. Image data were acquired in the Advanced Neural Imaging Center at KBRI. This research was supported by Brain Research Program (NRF-2017M3C7A1048089; J.W.K.) and Young Researcher Program (NRF-2020R1C1C1012788; T.-Y.C.) through the National Research Foundation of Korea (NRF), by KBRI basic research program through Korea Brain Research Institute (22-BR-04-03; J.W.K.) funded by Ministry of Science and ICT, and by Basic Science Research Program through the NRF funded by the Ministry of Education (NRF-2017R1A6A3A01076049; T.-Y.C.).

## Author contributions

Conceptualization & Methodology, T.-Y.C., H.J., M.C., and J.W.K.; Formal Analysis, T.-Y.C., H.J., S.J., and B.K.; Investigation, T.-Y.C., H.J., S.J., E.J.K., J.K., and Y.H.J.; Writing – Original Draft, T.-Y.C.; Writing – Review & Editing, T.-Y.C., H.J., S.J., J.K., M.C., and J.W.K.; Visualization, T.-Y.C.; Supervision, M.C. and J.W.K.; Funding Acquisition, T.-Y.C. and J.W.K.

## STAR Methods

### RESOURCE AVAILABILITY

#### Lead Contact

Further information and requests for resources and reagents should be directed to and will be fulfilled by the Lead Contact, Ja Wook Koo.

#### Materials Availability

This study did not generate new unique reagents.

#### Data and Code Availability

The data and code that support the findings from this study are available from the Lead Contact upon request. scRNA-seq data is deposited at NCBI’s Gene Expression Omnibus, and is publicly available through GEO Series accession number GSE222185.

### EXPERIMENTAL MODEL AND SUBJECT DETAILS

#### Animals

Adult (over 2 months old) male C57BL/6N mice and Fos-creER^T^^2^ mice were used for experiments. Mice of similar body weight within 10 percent differences and ages within a week of birth date were housed in cages of 2-4 for at least two weeks before surgical manipulations or behavioral experiments, maintained in an animal facility with a specific pathogen-free barrier under a 12-h light/dark cycle (lights on at 08:00 am), the temperature at 22 ± 2 ℃ and the humidity at 50 ± 10%, and given access to food and water *ad libitum*. All surgical manipulations and behavioral tests were carried out in the animal facility during their light on period. The Institutional Animal Care and Use Committee (IACUC) at Korea Brain Research Institute authorized all procedures (IACUC-21-00039).

### METHOD DETAILS

#### Stereotaxic surgeries

Mice were deeply anesthetized by an intraperitoneal injection of 0.1M phosphate-buffered saline (PBS) containing ketamine (100 mg/kg) and xylazine (10 mg/kg). The head of a deeply anesthetized mouse was fixed in a stereotaxic apparatus (Stoelting Co.), and the eyes were coated with ophthalmic ointment to avoid eye dryness. Scalp hair was removed, and the skin was cleaned with Betadine. Fine surgical scissors were used to make a midline incision to expose the skull, and a microdrill was used to make the craniotomies above the locations where viral vectors were injected or fiber optic cannulas were implanted. For all injections, a 5-μL microsyringe (7641-01, Hamilton) connected with a 33-gauge needle (7762-06, Hamilton) was used. Viral vectors were injected at a rate of 100 nL/min, and the needle was kept in place for an additional 10 min to allow for viral dispersion before being removed. The scalp incision was closed with a surgical suture and tissue adhesive (Vetbont, 3M). For implanting fiber optic cannulae, dental cement was applied to secure them. All stereotaxic coordinates were measured relative to bregma. After surgery, mice recovered in a clean cage under a heat lamp until awakening from anesthesia. Before undergoing subsequent operations or behavioral tests, mice were given a minimum of two weeks to recuperate.

For tracing activated mPFC neurons and projections (Figure 1A-E, S1A, B), Fos-creER^T^^2^ mice were bilaterally injected with 600 nL of a mixture of two viruses (AAV5-hSyn-mCherry and AAV5-EF1a-DIO-EYFP; ratio 1:1) into the mPFC (anteroposterior (AP), +2.1 mm; mediolateral (ML), ±0.6 mm; dorsoventral (DV), -2.1 mm; angle, 10°).

For TRACE experiments (Figure 1F-H), Fos-creER^T^^2^ mice were bilaterally injected 500 nL of AAVrg-EF1a-DIO-EYFP into the NAc (AP, +1.3 mm; ML, ±1.6 mm; DV, -4.6 mm; angle, 10°) and 500 nL of AAVrg-EF1a-DIO-mCherry into the VTA (AP, -3.2 mm; ML, ±1.2 mm; DV, -4.6 mm; angle, 10°).

For retrograde tracing experiments (Figure S1C-H, S2), C57BL/6N mice were bilaterally injected 500 nL of AAVrg-hSyn-EGFP into the NAc (AP, +1.3 mm; ML, ±1.6 mm; DV, -4.6 mm; angle, 10°) and/or 500 nL of AAVrg-hSyn-mCherry into the VTA (AP, -3.2 mm; ML, ±1.2 mm; DV, -4.6 mm; angle, 10°).

For genetic ablation experiments (Figure 2A-H, S2O, P), C57BL/6N mice were bilaterally injected 500 nL of AAVrg-EF1a-mCherry-IRES-Cre into the NAc (AP, +1.3 mm; ML, ±1.6 mm; DV, -4.6 mm; angle, 10°) or the VTA (AP, -3.2 mm; ML, ±1.2 mm; DV, -4.6 mm; angle, 10°) and 500 nL of AAV2-EF1a-Flex-taCasp3-TEVp or AAV2-EF1a-DIO-EYFP into the mPFC (AP, +2.1 mm; ML, ±0.6 mm; DV, -2.1 mm; angle, 10°).

For mPFC terminal activation experiments (Figure 2I-T, S2Q, R), C57BL/6N mice were unilaterally injected 500 nL of AAV5-CaMKIIa-hChR2(H134R)-EYFP or AAV5-CaMKIIa-EYFP into the right mPFC (AP, +2.1 mm; ML, ±0.3 mm; DV, -2.0 mm; angle, 0°), and fiber optic cannula (200 μm core diameter, 0.22 numerical aperture (NA); RWD Life Science) was implanted into the right NAc (AP, +1.3 mm; ML, +0.8 mm; DV, -4.3 mm; angle, 0°) or the right VTA (AP, -3.2 mm; ML, +0.4 mm; DV, -4.3 mm; angle, 0°) on the same day or several days after virus injection.

For fiber photometry experiments (Figure 3, S3), 500 nL of AAVrg-CAG-Cre was injected into the right NAc (AP, +1.3 mm; ML, +1.6 mm; DV, -4.6 mm; angle, 10°) or the right VTA (AP, -3.2 mm; ML, +1.2 mm; DV, -4.6 mm; angle, 10°) and 500 nL of AAVdj-DIO-GCaMP6s was injected into the right mPFC (AP, +2.1 mm; ML, +0.3 mm; DV, -2.0 mm; angle, 0°) of C57BL/6N mice. A fiber optic cannula (400 μm core diameter, 0.50 NA; RWD Life Science) was implanted 200 μm above the viral injection coordinates of the mPFC on the same day or several days after the virus injection.

For projection-specific genetic manipulation experiments (Figure 5, S5), C57BL/6N mice were bilaterally injected 500 nL of AAVrg-CAG-Cre into the VTA (AP, -3.2 mm; ML, ±1.2 mm; DV, -4.6 mm; angle, 10°) and 500 nL of AAV2-CMV-Flex-*Pou3f1* shRNA-EGFP-WPRE, AAV2-CMV-Flex-scrambled shRNA-EGFP, AAV2-CMV-Flex-*Pou3f1*-P2A-EGFP-WPRE, or AAV2-CAG-Flex-EGFP-WPRE into the mPFC (AP, +2.1 mm; ML, ±0.6 mm; DV, -2.1 mm; angle, 10°).

For mPFC-VTA cell body activation experiments (Figure 6, S6), C57BL/6N mice were bilaterally injected 500 nL of AAVrg-CAG-Cre into the VTA (AP, -3.2 mm; ML, ±1.2 mm; DV, -4.6 mm; angle, 10°) and 500 nL of AAV5-EF1a-DIO-hChR2(H134R)-EYFP or AAV5-EF1a-DIO-EYFP into the mPFC (AP, +2.1 mm; ML, ±0.6 mm; DV, -2.1 mm; angle, 10°). Fiber optic cannulae were bilaterally implanted 200 μm above the viral injection coordinates of the mPFC on the same day or several days after the virus injection.

#### Behavioral assays

##### Tube test

The tube test was conducted as instructed with minor modifications^20,22,66^. Mice were softly handled for at least 2 days (2 min/day) before the tube test to acclimate them to the experimenters and minimize stress and then trained to pass through the transparent acrylic tube with a 3-cm inner diameter and 30-cm length for 10 trials per day. On test days, two mice were released from the opposite sides of a tube, met at the center, and the mouse who withdrew first from the tube was declared as the “loser”. One more test with the same pair of mice was conducted after at least 1 minute. Mice were released at the opposite ends of the tube. It was considered a draw if the different mouse alternately wins in two repeated tests of the same pairs. For a set of 4 mice, the 6 pairs of mice were tested daily using a round-robin design in a random sequence, and the total number of wins on each test day was used to establish the social rank of the animals. If the same rank position in a cage did not change for the final two-thirds of the test time, this cage was used for further manipulation experiments.

For tracing activated mPFC neurons and projections (Figure 1A-E, S1A, B) or TRACE experiments (Figure 1F-H), 3 mice were housed in a cage for at least 2 weeks after virus injection and recovery. We conducted the same handling and tube training procedures (described above) for all mice. The tube test was repeated 10 times with a 1-min interval between the randomly selected pair of mice in a cage. The control mouse, which remained in the same cage, was permitted to cross the tube 10 times without competition with another mouse. Tamoxifen (T5648, Sigma-Aldrich; 150 mg/kg)^37,38^ or 4-hydroxytamoxifen (H6278, Sigma-Aldrich; 10 mg/kg)^40,41^ was intraperitoneally injected at 24 hours before or 2 hours after the tube test, respectively. After the behavioral tests, all mice were kept separately for a further 2 weeks to avoid the labelled neurons and projections activated by various social interactions under group housing and to fully express fluorophore.

For genetic ablation experiments (Figure 2A-H), a mouse that was injected viral vectors for projection-specific genetic ablation and another one that was injected viral vectors to express projection-specific fluorescent proteins were housed in a cage for at least 2 weeks after virus injection and recovery, and these 2 mice were conducted the same procedure of the tube test (described above) for 3 consecutive days to determine winner and loser.

For optogenetic activation (Figure 2I-T, S2Q, R, 6, S6) or fiber photometry experiments (Figure 3, S4), the same diameter and length tube with a 12 mm open slit on the top was used to pass the mice connected to the patchcords. Patchcords were connected to all mice at least 30 min before the training and test. For optogenetic activation experiments (Figure 2I-T, S2Q, R, 6, S6), the tube test ranks were verified once again before optogenetic stimulation, and blue light stimulation (473 nm, 1 ∼ 20 mW at the tip of the optical fiber; 10 ms pulses at 20 Hz for mPFC terminal activation, 10 ms pulses at 10 Hz for mPFC-VTA cell body activation) was delivered from right before mice entering the tube throughout the test. Light intensity was steadily increased until the rank changed or the maximal light intensity (20 mW) was achieved. For fiber photometry experiments (Figure 3, S3), a modified tube with a movable door at the middle of the tube was used. Mice in the tube started to compete when a door was removed after 10 s delay period to avoid any obstructions from the uncontrollable moving of mice in the tube or human handling.

With the exception of the tube test with fiber photometry recordings, all tube tests were videotaped using a webcam (C922, Logitech), and recordings of each tube test were blindly evaluated frame by frame. Five forms of behaviors during the tube test were manually annotated: 1) approach (either or both mice moved toward each other), 2) push (a mouse actively shoved the opponent), 3) resistance (a mouse held on the territory when being pushed by the opponent), 4) active retreat (a mouse voluntarily walked backward without pushes or approaches of the opponent) and 5) passive approach (a mouse backed away by pushes of the opponent). The number of each behavior during an entire tube test session was counted.

##### Wet bedding avoidance test

A rectangular plastic cage (20 cm X 35 cm) with new bedding was placed in the open field test box (40 cm X 40 cm) to block visual cues around the cage, and 250-300 mL of clean water was poured into the cage to fully dampen the bedding. A cylindrical acrylic platform (5 cm diameter and 5 cm height) was placed in the center of the cage. Four mice from the same cage were first entered into a cage with wet bedding, and all mice were briefly put on the platform for a while to make the mice recognize it and then lower them back to the floor. Behaviors of the four mice in the cage with wet bedding were videotaped for 20 min using a webcam (C922, Logitech). Each animal’s platform occupancy duration was then measured.

##### Direct social interaction test

This test was carried out with slight modifications as described in a prior study^67^. Mice were habituated at least 30 min prior in a behavior room under standard room light. A novel mouse (C57BL/6N, 5-6-week-old, male) was introduced in a home cage that was housing an experimental mouse. The total amount of social contacts of an experimental mouse, including anogenital and nose-to-nose sniffing, following, and allogrooming, was measured for 5 min.

#### Histology

Animals were deeply anesthetized with CO_2_ and transcardially perfused with 0.1 M phosphate-buffered saline (PBS) and 4% (wt/vol) paraformaldehyde (PFA) in PBS. Brains were removed, post-fixed overnight in 4% PFA, and then sectioned using a vibratome (VT1000S, Leica Biosystems). Or fixed brains were equilibrated in 30% sucrose in PBS at 4 °C and 50 or 60 μm thick coronal sections were cut on a cryocut microtome (CM1860, Leica Biosystems) at -20°C. All sections were washed in PBS prior to mounting on a slide glass or immunostaining.

To quantify activated mPFC neurons and their projections (Figure 1A-E, S1A, B), all brain sections were blocked with 0.3% Triton-X 100 and 3% bovine serum albumin in PBS for 1 h at room temperature, and then incubated in chicken anti-GFP antibody (1:500, GFP-1020, Aves Labs) and rat anti-RFP antibody (1:500, 5f8-100, ChromoTek) overnight. The following day, sections were incubated with donkey anti-chicken Alexa Fluor 488 antibody (1:500, 703-545-155, Jackson ImmunoResearch) and donkey anti-rat Alexa Fluor 594 antibody (1:500, A-21209, Invitrogen) for 3 hours. All sections were rinsed with PBS for 3 × 10 min and then mounted on slide glasses (Muto pure chemicals) and cover-slipped with VECTASIELD HardSet Antifade Mounting Medium with DAPI (H-1500, Vector Laboratories). Images were acquired using a fluorescence microscope (Pannoramic Scan system, 3DHISTECH) at the same gain and exposure with a 20X objective lens.

Immunostaining of c-fos and quantification (Figure S1C-H) were carried out as previously reported with minor modifications^68^. To count the number of c-fos-positive cells in the mPFC-NAc or mPFC-VTA (Figure S1C-H), the subject mice were sacrificed 90 min after performing the last tube test. Coronal sections containing the mPFC were prepared for c-fos immunostaining, and the viral injection sites were confirmed in sections containing the NAc and VTA. The mPFC sections were incubated in blocking solution (0.3% triton-X 100 and 4% normal donkey serum in PBS) for 1 h at room temperature and then incubated with rabbit anti-c-fos antibody (1:2,000, 2250S, Cell signaling technology) overnight at 4 °C. Following 3 × 10 min washing in PBS, incubation with donkey anti-rabbit Alexa Fluor 647 (1:200, A-31573, Invitrogen) was done for 3 h at room temperature. Sections were washed with PBS for 3 × 10 min, incubated in DAPI (1:5,000, 62248, Thermo Scientific) solution, and washed again. Stained sections were mounted on slides (Muto pure chemicals) and cover-slipped with VectaMount Permanent Mounting Medium (H-5000, Vector Laboratories). All images were obtained using ultra high-speed & spectral confocal microscopy (A1 Rsi/Ti-E, Nikon, Japan) at equal gain and exposure with a 10X objective lens.

For quantification of the efficiency of *Pou3f1* knockdown or overexpression (Figure S5AE-D), coronal sections containing the mPFC were immunostained for Pou3f1 by incubating with rabbit anti-Pou3f1 antibody (1:500, ab272925, abcam) overnight. The brain sections were then exposed to donkey anti-rabbit Alexa Fluor 555 antibody (1:500, A-31572, Invitrogen) for 2 hours. After washing and mounting on the slide slice, images were acquired using a fluorescence microscope (Pannoramic Scan system, 3DHISTECH) at the same gain and exposure with a 20X objective lens.

For histological experiments that do not require immunostaining (Figure 1G, 2B, 2F, 2J, 2P, 3C), steps after antibody treatment were performed in the same manner.

#### Fiber photometry

All fiber photometry experiments were conducted using Doric Fiber Photometry System (Doric Lenses). The two connectorized LEDs (CLEDs, 405 nm regulated at 208.616 Hz for calcium-independent signals and 465 nm regulated at 572.205 Hz for calcium-dependent signals) were controlled by the fiber photometry console via the LED driver. Each CLED was coupled to the seven-port Fluorescence MiniCube (FMC7_E1(400-410)_F1(420-450)_E2(460-490)_F2(500-540)_E3(555-570)_F3(580-680)_S) via an attenuating patch cord, and S port of the MiniCube was coupled to a pigtailed fiber optic rotary joint and a mono fiber-optic patchcord to deliver the excitation light to and to receive emitted light from behaving mice. The F2(50-540) port of the MiniCube was coupled to the photoreceiver (2151 Femtowatt Silicon Photoreceiver, Newport) or the photodetector (Fluorescence Detector Amplifier, Doric Lenses) that was coupled to an analog port of the fiber photometry console. Fiber photometry signals were recorded by Doric Neuroscience Studio (Ver. 5.3.3.14) through the Lock-In mode and a sampling rate of 12.0 kS/s. Tube tests with fiber photometry were recorded by a behavior tracking camera (Doric Lenses) controlled by the same software to synchronize the tube test behaviors and the recorded neural activities.

#### *Ex vivo* electrophysiology

All electrophysiological experiments were conducted 1 day after the final tube test, and brain slices from R1 and R4 mice were prepared at the same time to minimize the change in conditions due to the time difference when the two mice were sacrificed separately. Electrophysiological recordings were performed as we previously described with minor modifications^69,70,71^. Briefly, mice were anesthetized with isoflurane. After decapitation, the brains were removed rapidly and then submerged in an ice-cold, oxygenated (95% O_2_ and 5% CO_2_), low-Ca^2+^ / high-Mg^2+^ dissection buffer containing 2.5 mM KCl, 1.23 mM NaH_2_PO_4_, 26 mM NaHCO_3_, 11 mM dextrose, 0.5 mM CaCl_2_, 10 mM MgCl_2_, and 212.7 mM Sucrose. Coronal brain slices (300 μm) containing the mPFC and NAc or VTA (to check injection sites of retrograde viruses) were cut using a vibratome (VT1200S, Leica Biosystems). After that, the brain slices were put in a recovery chamber filled with oxygenated artificial cerebrospinal fluid (ACSF), which contained 124 mM NaCl, 2.5 mM KCl, 1.23 mM NaH_2_PO_4_, 26 mM NaHCO_3_, 11 mM dextrose, 2.0 mM CaCl_2_, and 1.0 mM MgCl_2_, and incubated at room temperature (24-26℃) for at least an hour before recording.

After recovery, the slices were moved to a submerged recording chamber where they were continuously perfused with oxygenated ACSF that was kept at room temperature at a flow rate of 2 ml/min. Slices were stabilized for at least 5 min prior to the recordings, and then whole-cell patch clamp recordings on retrogradely labelled mPFC pyramidal neurons were made using borosilicate glass pipettes (4-6 MΩ) filled with the internal solution containing 123 mM K-gluconate, 12 mM KCl, 10 mM HEPES, 0.2 mM EGTA, 4 mM Mg-ATP, 0.3 mM Na-GTP, and 10 mM Na_2_-phosphocreatine at pH 7.2-7.4 and 280-290 mOsm. Neuronal intrinsic properties such as excitability (Figure S2A, B, H, I), rheobase current measurement (Figure S2C, D, J, K), membrane capacitance (Cm) (Figure S2E, L), resting input resistance (R_In_) (Figure S2F, M), and resting membrane potential (RMP) (Figure S2G, N) were recorded under the current clamp mode. Only cells having an input resistance of more than 100 MΩ and an access resistance of less than 20 MΩ were recorded. The cells were excluded if the input or access resistance changed by more than 20%. Signals were amplified by a Multiclamp 700B (Molecular devices) and filtered at 2 kHz and digitized at 10 kHz with Digidata 1550B (Molecular Devices). Data were recorded and analyzed using pClamp 10 (Molecular Devices).

#### Tissue collection and library preparation for single-cell RNA sequencing

Animals were anesthetized with CO_2_ and the brains were extracted and transferred into ice-cold Dulbecco’s phosphate-buffered saline (DPBS) one day after the last tube test. For single cell dissociation, the brains were sectioned into 1.0 mm coronal slices in ice-cold DPBS with the brain matrix, and the mPFC was collected from each slice using 14-gauge punches (BP-10F, Kai Medical). In total, five pooled independent biological replicates including tissues from two-to-three different mice were analyzed for sequencing. Briefly, the tissues were cut into small pieces and incubated in a dissociation media (Hibernate A without Calcium and Magnesium with 1 mg/ml papain, 50% D-(+)-trehalose dihydrate, 25 mM DL-2-amino-5-phosphonopentanoic acid and DNase I) at 37 °C for 60 min with shaking at 225 RPM. After washing with 5 ml Hibernate A medium, the tissues were mechanically triturated with cutting pipette tips (tip diameter 2 mm and 1 mm) in 2 ml Hibernate A medium, which was repeated for another two times, and filtered through a cell strainer with 70 μm of the pore size to release single cells. The 1.5 ml single-cell suspension was pooled and loaded on a gradient medium (Hibernate A without Calcium and Magnesium with 1 mg/ml papain, 50% D-(+)-trehalose dehydrate, 25 mM DL-2-amino-5-phosphonopentanoic acid, DNase I, and ovomucoid inhibitor-albumin) and centrifuged at 900 RPM for 6 min at 4 °C to remove debris. The cells were then washed with 1 ml Hibernate A medium followed by 1 ml DPBS. The cells were spun down at 1,250 RPM for 6 min and re-suspended in DPBS. Countess II Automated Cell Counter (Invitrogen) was used to count the number of cells after centrifugation.1,000 ∼ 1,800 cells/μl or 1,500 ∼ 3,200 cells/μl were obtained from samples from two or three pooled mice, respectively.

For single cell RNA-sequencing (scRNA-seq), the cell suspension was diluted to the recommended concentration (700 ∼ 1,200 cells/μl) of the manufacturer’s protocol. We prepared cell suspension to 10,000 targeted cell recovery, and it was captured with 10X Chromium platform (10X Genomics, CA). 10x Genomics Chromium platform delivers a microfluidic chip for 3’ digital gene expression by profiling 500-10,000 individual cells per sample. Using Chromium Single Cell 3’ Reagent Kits v3, we prepared single cell master mix and gel beads, and then generated Gel Beads-in-emulsion (GEMs), combining barcoded Single Cell 3’ v3 Gel Beads. Next, the Gel Beads were dissolved, and cDNA was synthesized according to the sequence included in the GEMs. Since cDNA was synthesized, GEMs were broken and silane magnetic beads were used to purify the first-strand cDNA. And then, cDNA was amplified to generate sufficient mass for library for sequencing. Libraries were generated and sequenced from the cDNA and 10x Barcode containing the GEMs. The final libraries contained the P5 and P7 primers used in Illumina bridge amplification and the concentration was 1.7∼2.7ug.

#### Single-cell real-time quantitative PCR

Single cell RT-qPCR (scRT-qPCR) was carried out as previously described with minor modification^72,73,74,75^. RNA was isolated from retrogradely labelled mPFC-NAc or mPFC-VTA neurons from the coronal brain section containing mPFC 1 day after the final tube test. All glassware was oven-baked at 180℃ for at least 12 h, and all buffers and solutions were prepared using nuclease-free water (AM9932, Invitrogen) to minimize RNase contamination. Preparation of brain slices and whole-cell patch clamp recordings were performed as described in the *Ex Vivo* Electrophysiology section. Cells were aspirated with borosilicate glass pipettes (4-6 MΩ) filled with the same internal solution for electrophysiological recordings added 20 μg/ml glycogen and RNase inhibitor (0.16 Units/μL; SUPERaseIn RNase inhibitor (20 U/μL); AM2694, Invitrogen). After making a whole-cell configuration, intracellular contents including the nucleus were gently aspirated by applying negative pressure^75^, and the shape and the holding current of aspirated neurons were monitored to check the quality of the aspiration. RNA extraction from collected intracellular contents and reverse transcription were performed using a Sing Cell-to-CT^TM^ qRT-PCR kit (4458237, Invitrogen). In the pre-amplification and real-time PCR procedures, Taqman gene expression assays (*Nptx2*, Mm00479438_m1; *Nrn1*, Mm00467844_m1; *Cnih3*, Mm01319298_m1; *Ankrd33b*, Mm07298789_m1; *Mpv17*, Mm00485133_m1; *Nrip3*, Mm00508049_m1; *Pou3f1*, Mm00843534_s1; *Ly6a*, Mm00726565_s1; *Rn18s*, Mm03928990_g1; Thermo Fisher Scientific) were used. Amplification of cDNA was performed using LightCycler 480 Instrument II (Roche, Swiss), and then all reactions were analyzed using ΔΔCt method as previously described^76^ with 18S ribosomal RNA (*Rn18S*) as a normalized control.

### QUANTIFICATION AND STATISTICAL ANALYSIS

#### Quantification of activated neurons and projections

The number of EYFP-expressed cells and mCherry-expressed total infected cells, or the fluorescent intensity of EYFP-and mCherry-expressing projections in the NAc, LHA, MDT, BLA, PAG, and VTA was measured in the region of interest (ROI, 500 × 500 μm) using ImageJ (NIH, USA). Images were background subtracted and mean pixel intensity was measured for each projection region. The fraction of activated neurons or projections is calculated by dividing the number of EYFP-positive cells or the fluorescent intensity of EYFP-positive projections by the number of mCherry-positive cells or the fluorescent intensity of mCherry-positive projections. Cell numbers or fluorescent intensity of projections were averaged from eight coronal sections per mouse in all quantifications. All procedures were done by a blinder observer.

#### Quantification of TRACE-labelled or c-fos-positive cells

The number of TRACE-labelled or c-fos-positive cells in the ROI (500 × 500 μm) was counted by ImageJ by a blinder observer. Cell numbers of eight coronal sections per mouse were averaged.

#### Fiber photometry

Data analysis was conducted using an open-source analysis package, named photometry modular analysis (pMAT)^77^. The onsets of each behavior were synchronized to time zero, and Z-score standardization was applied to the signals. The averaged response to each tube test behavior was calculated by subtracting the average Z-scored data 1 s prior to and 2 s after the behavioral onset. The area under the curve (AUC) per second values were calculated to compare the neural activity before and after the behavioral onset.

#### scRNA-seq data processing, clustering, and annotation

Raw sequencing data from each sample were quantified using 10x Genomics Cell Ranger 4.0.0^78^. Mouse mm10 reference recommended and provided by 10x Genomics (Pleasanton, CA) was used. Then, Seurat package 3.1.3 in R was used for subsequent data processing^79^. As a quality control, cells that expressed too few or too many genes were excluded (Table S1). In addition, cells with high expression rates of mitochondrial, ribosomal, and hemoglobin genes were considered low-quality and excluded. Details on filtering criteria and the number of cells generated from each sample are in Table S1. Then, Log Normalization and data integration to correct batch effect were conducted using Seurat. For clustering, the first 20 principal components were used to create k-nearest neighbor (KNN) graphs and Uniform Manifold Approximation and Projection (UMAP) and FindClusters function, implementing the Louvain algorithm, in Seurat with the resolution of 0.2 was used. Cell type annotation was performed using the SingleR package^80^. In-house cluster reference gene lists were used for annotation.

#### mPFC-NAc or mPFC-VTA classification and DEG analysis

Neuronal cells were isolated and subjected to PCA, UMAP, and clustering based on the expression of enriched genes of NAc-or VTA-projecting mPFC neurons in the previously reported study^28^ using the same parameter. Seurat’s FindMarkers function was used to find the marker gene for each cluster with logfc.threshold parameter of 0.25, the test method of the Wilcoxon Rank Sum test. Genes with p-adj < 0.05 was defined as cluster marker genes. Then, DEGs were found between R1 and R4 using the FindMarkers function with the same parameters.

#### Pou3f1 chromatin immunoprecipitation sequencing data analysis

We downloaded Pou3f1 chromatin immunoprecipitation sequencing (ChIP-seq) peak bed files from previous studies (corresponding GEO IDs: GES110950, GSE74636, GSE54569, and GSE69778). ChIPseeker package 1.32.1 was used to annotate peak. Transcription start site regions were defined as ± 3 kb of transcription start sites. TxDb.Mmusculus.UCSC.mm9.knownGene R package was used as an annotation database. Gene ontology (GO) enrichment analysis was conducted using clusterProfiler package 4.4.4 with multiple test-adjusted P-value threshold of 0.05.

#### scRT-qPCR and clustering

The fold change values (2^-ΔΔCT^) of cluster markers (C8, *Ly6a*; C0, *Nrip3*) were used to cluster C1 or C8 from mPFC-NAc neurons or C0 or C3 from mPFC-VTA neurons. The Expectation-Maximization (EM) or Simple K-means analysis in Machine Learning Algorithm in JAVA (WEKA, New Zealand) system was used for the clustering of the scRT-qPCR data. The mRNA level of DEGs was compared according to the clustering results.

#### Statistical analysis

Every experiment used anonymous samples, and the experimenters were not informed of the experimental conditions of the animals. Statistical analyses were conducted using GraphPad Prism 9 and are presented as mean ± the standard error of the mean (SEM). Comparisons between the two groups were determined by Student’s t-test, Mann-Whitney U test, Chi-square test, Wilcoxon Rank-Sum test, or Wilcoxon matched-pairs signed rank test. Multiple comparisons were determined by one-way analysis of variation (ANOVA) or Kruskal-Wallis test followed by post-hoc testing. The significance criterion was set at a P-value of ≤0.05. Comprehensive information on statistical analysis is included in the figures, figure legends, and Table S3.

## Supplemental information titles and legends

**Figure S1.**
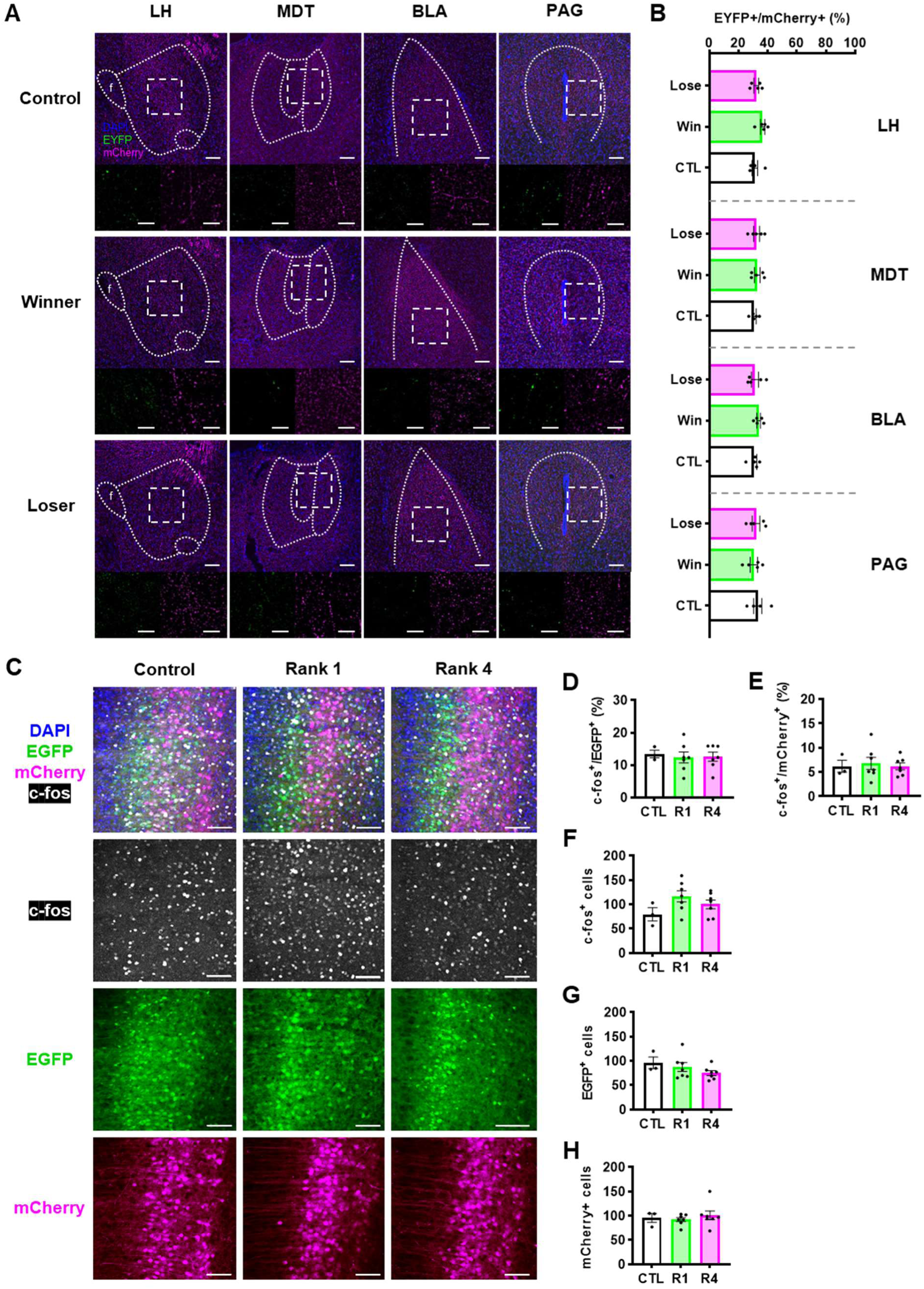
Neuronal activity of the mPFC-NAc or mPFC-VTA between social dominants and subordinates, related to Figure 1. (A) Representative images containing images of projections in the LHA, MDT, BLA and PAG and magnified images of the area indicated by a white dashed box. Scale bars, 0.1 mm (top); 0.01 mm (bottom). (B) Quantification of activated projections by the ratio of EYFP^+^ (or FosTRAPed) axon terminals per mCherry^+^ axon terminals (or total infected mPFC terminals) in ROIs. n = 5 mice in each group; P = 0.0944 (LHA), P = 0.7047 (MDT), P = 0.5090 (BLA), and P = 0.7679 (PAG) by one-way ANOVA with Tukey’s test. (C) Representative images of the mPFC from control (left), rank 1 (R1, center) or rank 4 (R4, right). Scale bars, 0.1 mm. (D, E) Quantitative analysis of c-fos^+^ neurons per EGFP^+^ (or mPFC-NAc) (D) or mCherry^+^ (or mPFC-VTA) (E) neurons in the mPFC. n = 3 mice (CTL), n = 7 (R1), and n = 7 (R4); P = 0.9176 (D) and P = 0.8967 (E) by one-way ANOVA with Tukey’s test. (F-H) Quantitative analysis of c-fos^+^ (F), EGFP^+^ (or mPFC-NAc) (G), or mCherry^+^ (or mPFC-VTA) (H) neurons in the mPFC. n = 3 mice (CTL), n = 7 (R1), and n = 7 (R4); P = 0.1667 (F), P = 0.2787 (G), and P = 0.6823 (H) by one-way ANOVA with Tukey’s test.

**Figure S2.**
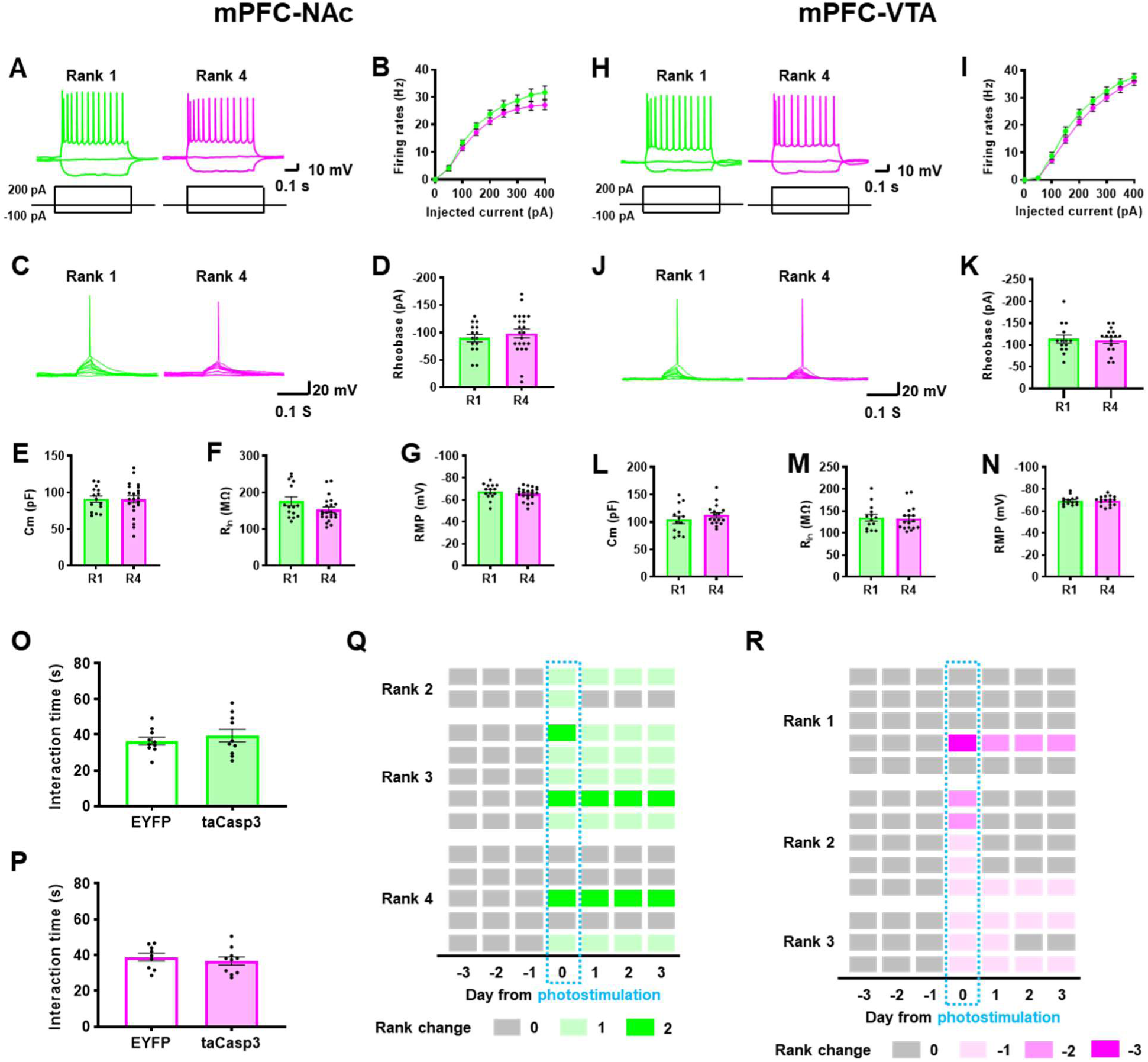
Intrinsic electrophysiological properties of the mPFC-NAc or mPFC-VTA neurons between social dominants and subordinates, related to Figure 1, and behavioral effects of manipulating mPFC-NAc or mPFC-VTA, related to Figure 2. (A, H) Representative traces of intrinsic excitability measured as firing rates in response to step depolarizing currents (50 pA increment, 500 ms duration) in mPFC-NAc (A) or mPFC-VTA (H) neurons. Left, rank 1 (R1); Right, rank 4 (R4). (B, I) Summary graphs of intrinsic excitability in mPFC-NAc (B) or mPFC-VTA (I) neurons. n = 15 neurons from 2 mice (A, R1), n = 22, 2 (A, R4), n = 14, 2 (H, R1), and n = 17, 2 (H, R4); P = 0.8325 (B, 50 pA), P = 0.2097 (B, 100 pA), P = 0.1806 (B, 150 pA), P = 0.1652 (B, 200 pA), P = 0.2125 (B, 250 pA), P = 0.2129 (B, 300 pA), P = 0.1083 (B, 350 pA), P = 0.9882 (B, 400 pA), P = 0.9344 (I, 50 pA), P = 0.2511 (I, 350 pA), or P = 0.2229 (I, 400 pA) by Mann-Whitney U test (two-tailed), P = 0.6518 (I, 100 pA), P = 0.2430 (I, 150 pA), P = 0.1597 (I, 200 pA), P = 0.1883 (I, 250 pA), and P = 0.2889 (I, 300 pA) by Student’s t-test (unpaired, two-tailed). (C, J) Representative traces of rheobase current measurement by brief step depolarizing currents (10 pA increment, 10 ms duration) in mPFC-NAc (A) or mPFC-VTA (H) neurons. Left, R1; Right, R4. (D, I) Summary graphs of rheobase current measurement in mPFC-NAc (A) or mPFC-VTA (H) neurons. n = 15 neurons from 2 mice (C, R1), n = 22, 2 (C, R4), n = 14, 2 (J, R1), and n = 17, 2 (J, R4); P = 0.4869 (D), and P = 0.7950 (I). (E, L) Comparison of the membrane capacitance (Cm) of mPFC-NAc (E) or mPFC-VTA (L) neurons. n = 15 neurons from 2 mice (E, R1), n = 22, 2 (E, R4), n = 14, 2 (L, R1), and n = 17, 2 (L, R4); P= 0.9882 (E) and P = 0.3017 (L) by Student’s t-test (unpaired, two-tailed). (F, M) Comparison of the resting input resistance (R_In_) of mPFC-NAc (F) or mPFC-VTA (M) neurons. n = 15 neurons from 2 mice (F, R1), n = 22, 2 (F, R4), n = 14, 2 (M, R1), and n = 17, 2 (M, R4); P= 0.2019 (F) or 0.9221 (M) by Mann-Whitney U test (two-tailed). (G, N) Comparison of the resting membrane potential (RMP) of mPFC-NAc (G) or mPFC-VTA (N) neurons. n = 15 neurons from 2 mice (G, R1), n = 22, 2 (G, R4), n = 14, 2 (N, R1), and n = 17, 2 (N, R4); P = 0.2866 (G) and P = 0.9708 (N) by Student’s t-test (unpaired, two-tailed). (O, P) Statistics for social interaction time of control versus mice inhibiting mPFC-NAc (A) or mPFC-VTA (B). n = 10 mice (A, EYFP in mPFC-NAc), n = 10 (A, taCasp3 in mPFC-NAc), n = 9 (B, EYFP in mPFC-VTA), and n = 10 (B, taCasp3 in mPFC-VTA); P = 0.4740 (A) and P = 0.5039 (B) by Student’s t-test (unpaired, two-tailed). (Q, R) Summary rank change of each manipulated mouse with optogenetic activation of mPFC-NAc (C) or mPFC-VTA (D). n = 12 cages (C), and n = 13 (D).

**Figure S3.**
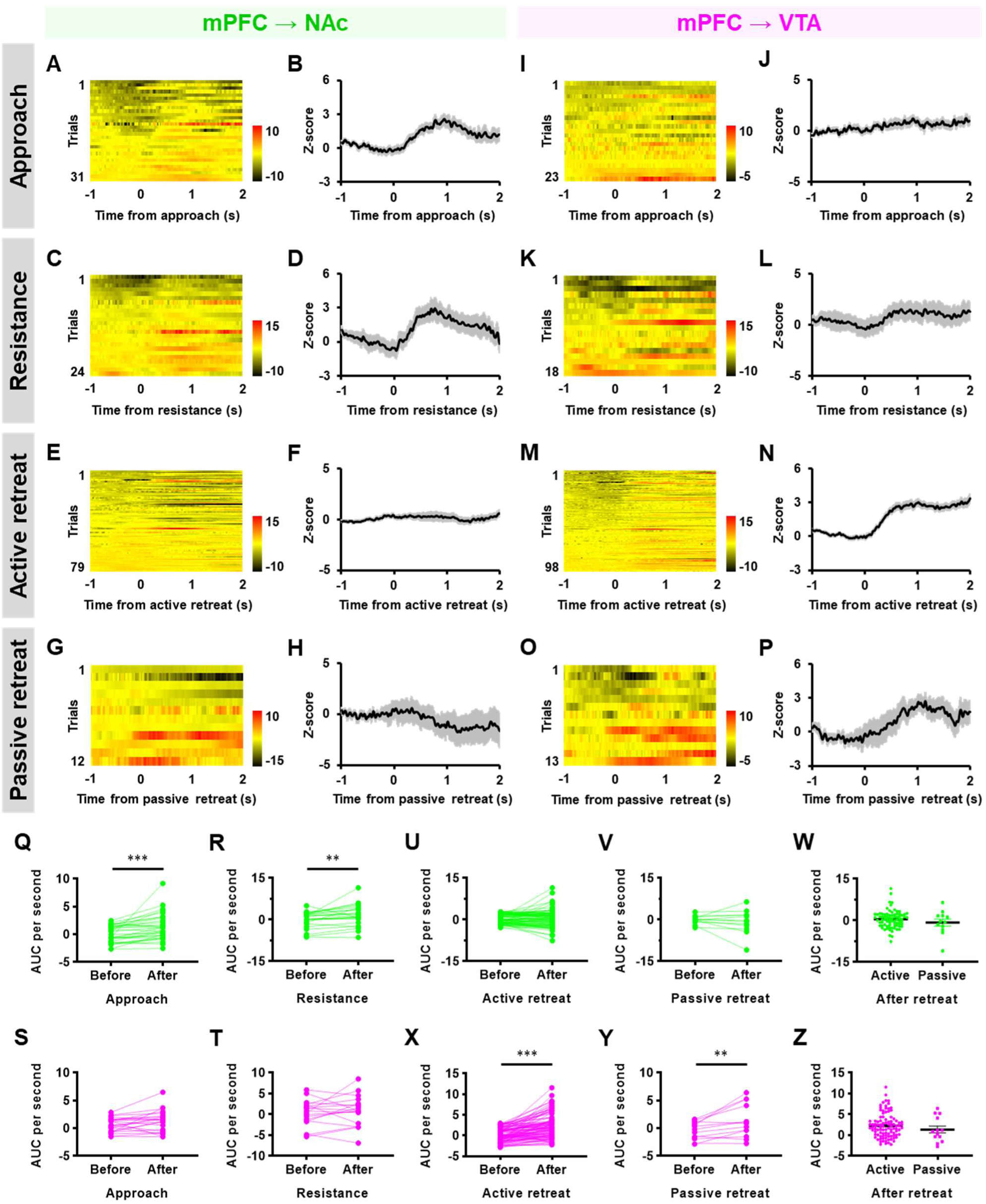
Other social competition behaviors were separately encoded in mPFC-NAc or mPFC-VTA, related to Figure 3. (A-H) Heatmap (A, C, E, G) and peri-event plot (B, D, F, H) of Z-scored mPFC-NAc activity aligned to the onset of approaches (A, B), resistances (C, D), active retreats (E, F) or passive retreats (F, G). n = 31 approach trials (A, B), n = 24 resistance trials (C, D), n = 79 active retreat trials (E, F), and n = 12 passive retreat trials (G, H) from 10 mice. (I-P) Heatmap (I, K, M, O) and peri-event plot (J, L, N, P) of Z-scored mPFC-VTA activity aligned to the onset of approaches (I, J), resistances (K, L), active retreats (M, N) or passive retreats (O, P). n = 31 approach trials (I, J), n = 18 resistance trials (K, L), n = 98 active retreat trials (M, N), and n = 13 passive retreat trials (O, P) from 10 mice. (Q-T) Quantification of the area under curve (AUC) per second before and after approaches (Q, S) and resistances (R, T) in mPFC-NAc (Q, R) or mPFC-VTA (S, T). n = 31 trials (Q), n = 24 (R), n = 23 (S), and n = 18 (T); P < 0.001*** (Q), P = 0.0598 (S), and P = 0.3624 (T) by Student’s t-test (paired, two-tailed) and P = 0.0016** (R) by Wilcoxon matched-pairs signed rank test (two-tailed). (U, V, X, Y) Quantification of the area under curve (AUC) per second before and after active retreat (U, X) and passive retreat (V, Y) in mPFC-NAc (U, V) or mPFC-VTA (X, Y). n = 79 trials (U), n = 12 (V), n = 98 (X), and n = 13 (Y); P = 0.2963 (U) and P < 0.001*** (V) by Wilcoxon matched-pairs signed rank test (two-tailed), and P = 0.4874 (X) and P = 0.0068** (Y) by Student’s t-test (paired, two-tailed). (W, Z) Comparison of the area under curve (AUC) per second between active and passive retreat of mPFC-NAc (W) or mPFC-VTA (Z). n = 79 active retreat and 12 passive retreat trials (W), and n = 98 active and 13 passive retreat trials (Z); P = 0.4063 (W) and P = 0.2000 (Z).

**Figure S4.**
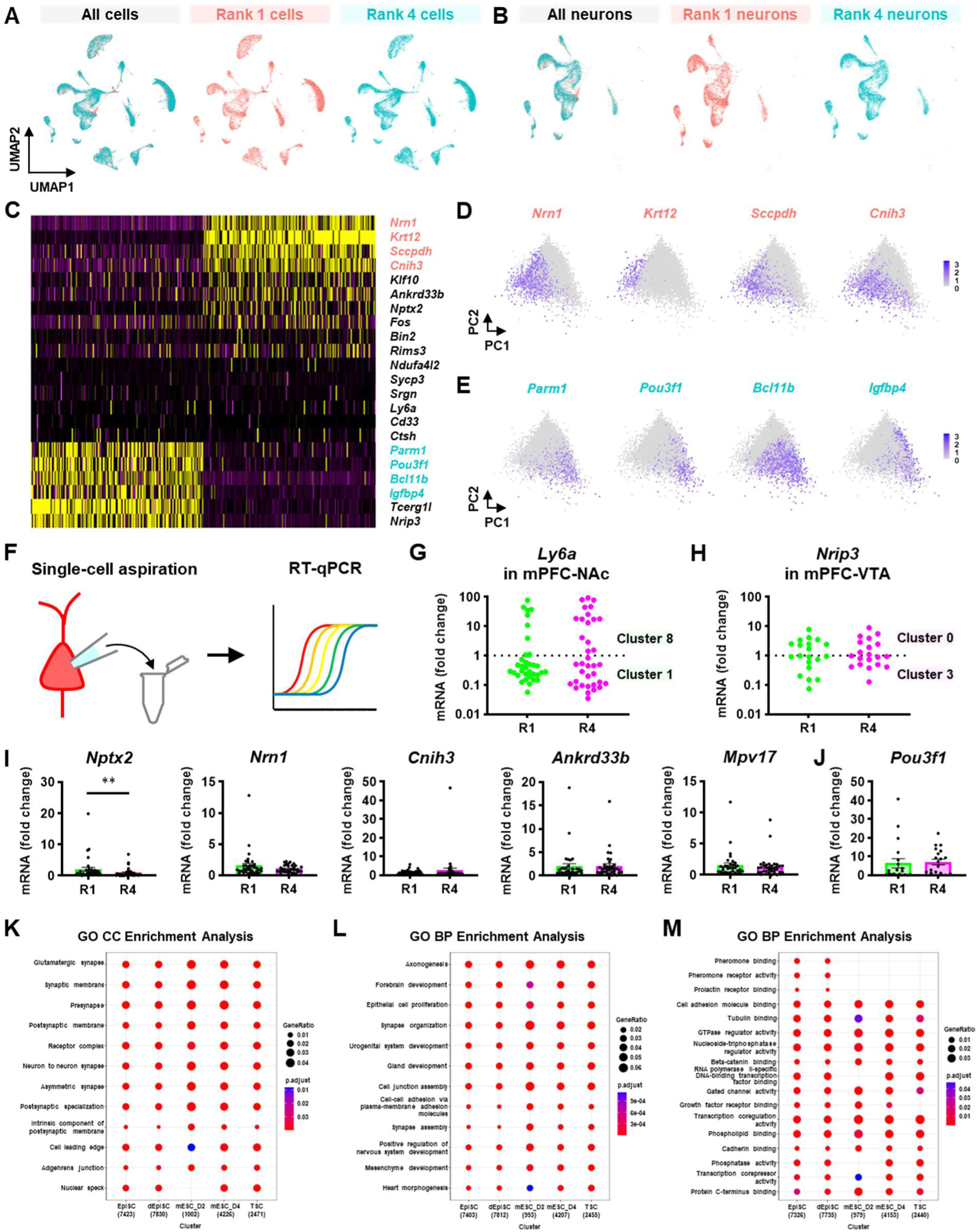
Projection-specific DEG analysis by scRNA-seq and validation by scRT-qPCR, related to Figure 4. (A) Two-dimensional UMAP visualization showing the distribution of all cells (left), cells from rank 1 (center) or 4 (right) among major mPFC cell types. (B) Two-dimensional UMAP visualization showing the distribution of all neurons (left), neurons from rank 1 (center) or 4 (right) among major mPFC neuronal subtypes. (C) Classification of NAc-or VTA-projecting mPFC neuronal subpopulations by principal component analysis (PCA) using projection-specific marker genes. (D, E) Distribution of top 4 marker genes of mPFC-NAc (D) or mPFC-VTA (E) in neuron-specific PCA plots. (F) Schematic illustration of scRT-qPCR of aspirated intracellular contents from retrogradely labelled mPFC neurons. (G, H) Clustering mPFC-NAc (G) or mPFC-VTA (H) by the expression level of cluster marker gene (i.e., *Ly6a* for C8 of mPFC-NAc; *Nrip3* for C0 of mPFC-VTA). n = 34 cells from 4 mice (G, R1), 36, 4 (G, R4), 20, 3 (H, R1), and 20, 3 (H, R4); P = 0.8221 (G) and P = 0.9201 (H) by Mann-Whitney U test (two-tailed). (I, J) scRT-qPCR validation of DEGs between R1 and R4 in mPFC-NAc (C1 + C8) (I) or mPFC-VTA (C0 + C3) (J). n = 34-35 cells from 4 mice (I, R1), 36, 4 (I, R4), 20, 3 (J, R1), oand 20, 3 (J, R4); P = 0.0036** (I, *Nptx2*), P = 0.5249 (I, *Nrn1*), P = 0.7195 (I, *Cnih3*), P = 0.2047 (I, *Ankrd33b*), P = 0.4960 (I, *Mpv17*), and P = 0.2395 (J, *Pou3f1*). (K-M) Gene ontology (GO) enrichment analysis of chromatin immunoprecipitation sequencing (ChIP-seq) of Pou3f1. Cellular component (CC) (K), biological process (BP) (L), and molecular function (MF) (M). Each circle is colored by multiple test-adjusted P-value and sized by the ratio of genes that have the GO term. EpiSC, epiblast stem cell; dEpiSC, differentiated epiblast stem cell; mESC, mouse embryonic stem cell; TSC, trophoblast stem cell.

**Figure S5.**
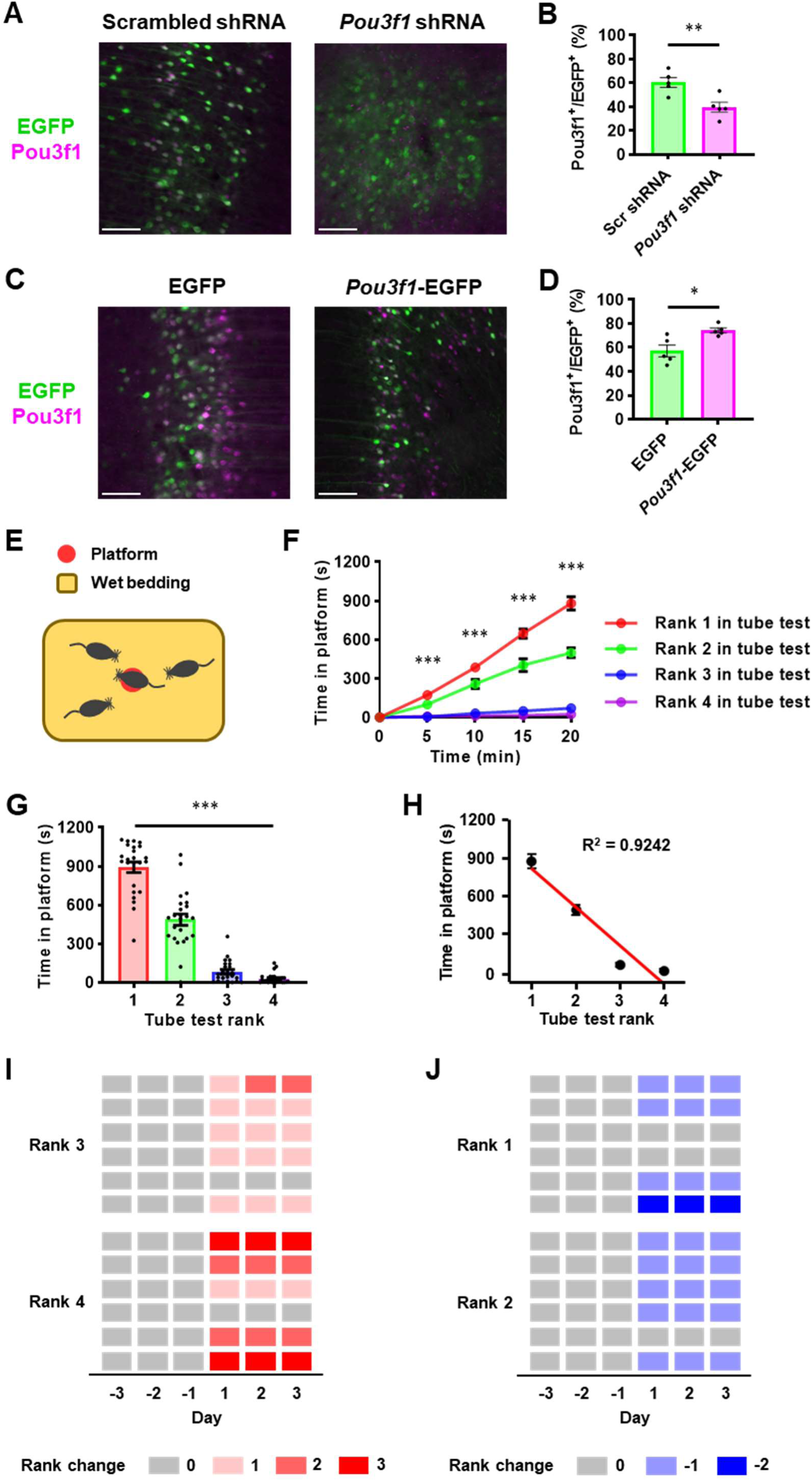
Behavioral effects of manipulating *Pou3f1* expression in the mPFC-VTA, related to Figure 5. (A) Representative images containing the mPFC infected by AAV2-Flex-scrambled shRNA-EGFP (left) or AAV2-Flex-*Pou3f1* shRNA-EGFP (right). Cre recombinase is retrogradely transported by injecting AAVrg-CAG-Cre in the VTA. Scale bars, 100 μm. (C) Representative images containing the mPFC infected by AAV2-DIO-EGFP (left) or AAV2-DIO-*Pou3f1*-EGFP (right). Cre recombinase is retrogradely transported by injecting AAVrg-CAG-Cre in the VTA. Scale bars, 100 μm. (B, D) Quantitative analysis of Pou3f1-expressing neurons per EGFP^+^ (or virus-infected mPFC-VTA) neurons in the mPFC. n = 5 mice (F, Scr shRNA), 5 (F, *Pou3f1* shRNA), 5 (H, EGFP), and 5 (H, *Pou3f1*-EGFP); P = 0.0071** (F) and P = 0.0110* (H) by Student’s t-test (unpaired, two-tailed). (E) Schematic illustration of the wet-bedding avoidance (WBA) test. Four cagemate mice compete for an elevated platform in the center of a cage to avoid wet bedding. (F-H) Cumulative time on the platform of four mice of different tube test rank (B). Total time on the platform in the 20-mintest (C). Correlation between time on the platform and tube test rank (D). n = 24 cages; P < 0.001*** (B, 5-20 min; C, Time in platform) by Friedman test with Dunn’ test. (I, J) Summary rank change of each manipulated mouse with *Pou3f1* K/D (I) or O/E (J). n = 12 cages (I), and n = 12 (J).

**Figure S6.**
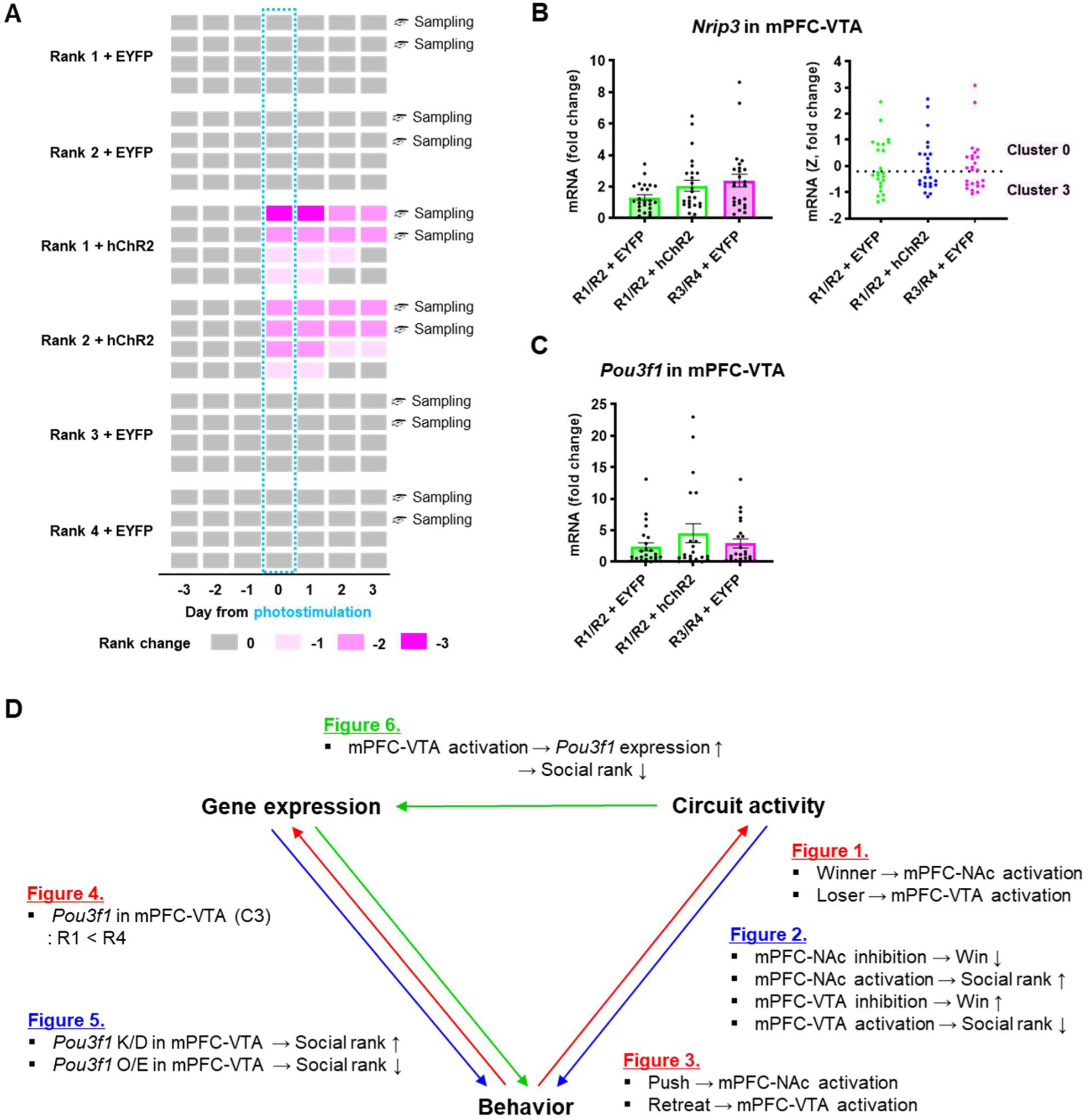
The effects of soma-targeting mPFC-VTA optogenetic activation on social dominance behavior and *Pou3f1* expression, related to Figure 6. (A) Summary rank change of each manipulated mouse with soma-targeting optogenetic activation of the mPFC-VTA. n = 8 cages (R1/R2 + EYFP), 8 (R1/R2 + hChR2), and 8 (R3/R4 + EYFP). (B) Clustering mPFC-VTA by the expression level of C0 marker gene *Nrip3*. *Nrip3* expression was not statistically different, but the average or the distribution is different between three groups (left). Therefore, *Nrip3* expression was converted to the standard deviation (Z-score) and then clustered because it induced incorrect clustering (right). n = 24 cells from 4 mice (R1/R2 + EYFP), n = 21, 4 (R1/R2 + hChR2), and n = 23, 4 (R3/R4 + EYFP); P = 0.0883 (fold change) and P = 0.9697 (Z, fold change) by Kruskal-Wallis test with Dunn’s test. (C) The expression level of *Pou3f1* mRNA between R1/R2 + EYFP, R1/R2 + hChR2, and R3/R4 + EYFP in mPFC-VTA (C0 + C4). n = 24 cells from 4 mice (R1/R2 + EYFP), n = 21, 4 (R1/R2 + hChR2), and n = 23, 4 (R3/R4 + EYFP); P = 0.9044 by Kruskal-Wallis test with Dunn’s test. (D) Summary of all findings in this study to reveal a causal relationship between circuit activity, gene expression and behavior

**Table S1.** Summary of cell filtering process, related to Figure 4.

| Sample |  | Filtering criteria |  |  |  | # of cells |  |
| --- | --- | --- | --- | --- | --- | --- | --- |
|  |  | Gene | Mt % | Ribo % | Hb % | Raw | Filtered |
| Rank 1 | Sample 1 | 500<nGene<6000 | <10% | <10% | <10% | 6,126 | 3,264 |
|  | Sample 2 | 200<nGene<4500 | <10% | <10% | <10% | 2,551 | 2,075 |
|  | Sample 3 | 200<nGene<4500 | <10% | <10% | <10% | 1,838 | 1,464 |
|  | Sample 4 | 300<nGene<6000 | <10% | <10% | <10% | 3,454 | 1,672 |
|  | Sample 5 | 400<nGene<5000 | <10% | <10% | <10% | 9,483 | 6,797 |
| Rank 4 | Sample 1 | 700<nGene<6500 | <10% | <10% | <10% | 10,103 | 1,426 |
|  | Sample 2 | 200<nGene<4500 | <10% | <10% | <10% | 2,321 | 2,011 |
|  | Sample 3 | 200<nGene<4500 | <10% | <10% | <10% | 4,630 | 3,841 |
|  | Sample 4 | 350<nGene<4000 | <10% | <10% | <10% | 8,605 | 5,438 |
|  | Sample 5 | 350<nGene<4000 | <10% | <10% | <10% | 15,826 | 13,421 |

● nGene: the number of genes expressed in a cell ● Mt %: the percentage of mitochondrial genes in a cell ● Ribo %: the percentage of ribosomal genes in a cell ● Hb %: the percentage of hemoglobin genes in a cell

**Table S2.** Summary of annotation, cell numbers and ratio of clustered cell types, related to Figure 4.

| Annotation | Rank 1 |  | Rank 4 |  | Total |  |
| --- | --- | --- | --- | --- | --- | --- |
|  | Cell number | Percent (%) | Cell number | Percent (%) | Cell number | Percent (%) |
| Neurons | 5325 | 34.87 | 9558 | 36.57 | 14883 | 35.94 |
| Astrocytes | 3651 | 23.91 | 5637 | 21.57 | 9288 | 22.43 |
| Oligodendrocytes | 1483 | 9.71 | 3003 | 11.49 | 4486 | 10.83 |
| Endothelial cells | 1459 | 9.55 | 2655 | 10.16 | 4114 | 9.94 |
| Microglia | 1783 | 11.67 | 2635 | 10.08 | 4418 | 10.67 |
| Fibroblasts | 794 | 5.20 | 1492 | 5.71 | 2286 | 5.52 |
| Epithelial cells | 212 | 1.39 | 321 | 1.23 | 533 | 1.29 |
| Macrophages | 295 | 1.93 | 97 | 0.37 | 392 | 0.95 |
| Etc. | 270 | 1.77 | 739 | 2.83 | 1009 | 2.44 |
| Total | 15272 | 100 | 26137 | 100 | 41409 | 100 |

**Table S3.** Summary of statistical analyses, related to Figure 1, 2, 3, 4, 5, and 6.

**Video S1.** Tube test with optogenetic activation of mPFC-NAc, related to Figure 2.

**Video S2.** Tube test with optogenetic activation of mPFC-VTA, related to Figure 2.

**Video S3.** Wet bedding avoidance (WBA) test, related to Figure 5.

